# Stress adaptation of mitochondrial protein import by OMA1-mediated degradation of DNAJC15

**DOI:** 10.1101/2025.03.04.641455

**Authors:** Lara Kroczek, Hendrik Nolte, Yvonne Lasarzewski, Thibaut Molinié, Daniel Curbelo Pinero, Kathrin Lemke, Elena Rugarli, Thomas Langer

**Author notes:** Corresponding author. Max Planck Institute for Biology of Ageing, Joseph-Stelzmann-Str. 9b, 50931 Cologne, Germany. These authors contributed equally (Lara Kroczek, Hendrik Nolte).

## Abstract

Mitochondria adapt to cellular stress to ensure cell survival. The stress-regulated mitochondrial peptidase OMA1 orchestrates these adaptive responses, which limit mitochondrial fusion and promote mitochondrial stress signaling and metabolic rewiring. Here, we show that cellular stress adaptation involves OMA1-mediated regulation of mitochondrial protein import and OXPHOS biogenesis. OMA1 cleaves the mitochondrial chaperone DNAJC15 and promotes its degradation by the m-AAA protease AFG3L2. Loss of DNAJC15 reduces the import of OXPHOS-related proteins via the TIMM23-TIMM17A protein translocase, limiting OXPHOS biogenesis under conditions of mitochondrial dysfunction. Non-imported mitochondrial preproteins accumulate at the endoplasmic reticulum and induce an ATF6-related unfolded protein response. Our results demonstrate stress-dependent changes in protein import specificity as part of the OMA1-mediated mitochondrial stress response and highlight the interdependence of proteostasis regulation between different organelles.

## Introduction

Mitochondrial stress responses orchestrated by the mitochondrial peptidase OMA1 ensure the adaptation of mitochondrial functions to meet changing physiological demands and maintain cell survival. Mitochondrial deficiencies, such as OXPHOS defects or dissipation of the membrane potential, activate OMA1 in the mitochondrial inner membrane (IM) and trigger the mitochondrial integrated stress response (ISR^mt^) by processing of DELE1, which promotes metabolic reprogramming and enhances the cellular defence against oxidative stress ^1–6^. At the same time, OMA1 activation limits mitochondrial fusion through excessive processing of the dynamin-like OPA1 leading to the fragmentation of the mitochondrial network under stress ^7–9^ ^10^. Recent studies have demonstrated that OMA1 can also perform quality control functions by degrading polypeptides that block protein translocases ^11^. The activation of OMA1 destabilizes the protease and causes its degradation, terminating the OMA1-mediated stress response after restoration of mitochondrial functions ^6,12^. Although impairment of OMA1-mediated stress pathways by *Dele1* deletion or prevention of Opa1 processing is well tolerated in mice ^13^, protective effects have been observed under pathophysiological conditions, such as mitochondrial (cardio-) myopathies ^14–17^. However, the consequences of OMA1 deficiency and of impaired OMA1-mediated stress responses are cell and tissue specific and depend on the physiological context ^18–23^.

Here, to better understand how OMA1 affects mitochondrial proteostasis and stress responses, we performed a proteomic survey for proteolytic substrates of OMA1. We demonstrate that OMA1 cleaves the mitochondrial chaperone DNAJC15 facilitating its degradation by the mitochondrial m-AAA protease. The loss of DNAJC15 alters protein import by TIM23 protein translocases in the IM and limits the accumulation of OXPHOS-related mitochondrial matrix and IM proteins. Non-imported mitochondrial preproteins accumulate at the endoplasmic reticulum (ER) and trigger an ATF6-related unfolded protein response. These results demonstrate that OMA1 allows to adapt mitochondrial protein biogenesis to stress and reveal an intricate network of cellular stress responses to proteostasis disturbances.

## Results

### OMA1 cleavage facilitates DNAJC15 degradation by the m-AAA protease

To identify novel OMA1 substrate proteins, we searched for amino-terminal peptides of mitochondrial proteins that accumulate in an OMA1-dependent manner in a proteome-wide survey in wild-type (WT) and *OMA1^−/−^* human cervical cancer cells (Fig. 1a, Supplementary Table 1). Among the four MitoCarta 3.0 annotated peptides ^24^, that were significantly decreased in *OMA1*^−/−^ cells, was one amino-terminal peptide of the protein DNAJC15 (MCJ), a cochaperone of the human TIM23 translocase in the mitochondrial inner membrane (IM) ^25–27^ (Extended Data Fig. 1a). We detected a peptide corresponding to amino acids 20-35, indicating OMA1 cleavage of DNAJC15 (Q9Y5T4) after amino acid 19.

**Fig. 1.**
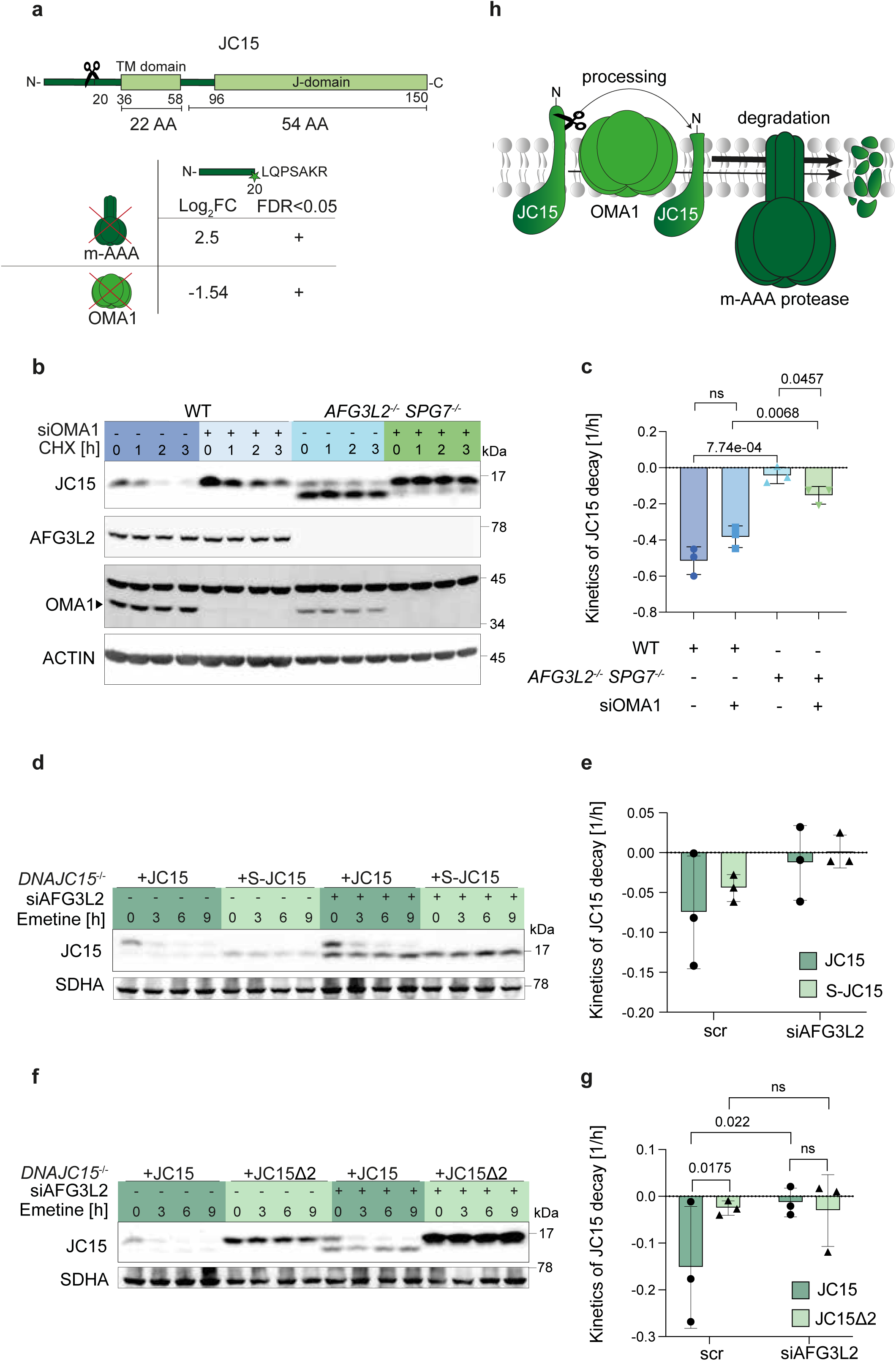
OMA1 cleavage of DNAJC15 facilitates degradation by the m-AAA protease. **a**, Comparison of the Neo-N-terminal proteome data sets of wild-type (WT) HeLa cells with *OMA1^−/−^* and *AFG3L2^−/−^ SPG7^−/−^* cells reveals the OMA1-dependent cleavage of DNAJC15 (JC15) at amino acid position 19 (UniProt ID: Q9Y5T4) (n = 5). **b**, Stability of DNAJC15 in WT and *AFG3L2^−/−^ SPG7^−/−^* HeLa cells depleted of OMA1 (siOMA1) for 72 h after inhibition of cytosolic translation with Cycloheximide (CHX). A representative experiment is shown (n = 3). **c**, Degradation rates of DNAJC15 in (b) normalized to time point 0 h. Data are means +/− SD (n = 3). *P* values were calculated using a two-sided unpaired t-test. **d**, Stability of DNAJC15 in *DNAJC15*^−/−^ cells expressing DNAJC15 (JC15) or DNAJC15 cleaved by OMA1 (S-JC15) and depleted of AFG3L2 (siAFG3L2) after inhibition of cytosolic translation with emetine. **e**, Degradation rates of DNAJC15 in (d) normalized to time point 0 h. Data are means +/− SD (n = 3). Statistics were calculated with a 2-Way ANOVA test. **f**, Stability of DNAJC15 in *DNAJC15*^−/−^ cells expressing DNAJC15 (JC15) or a non-cleavable DNAJC15 variant lacking amino acid 19 and 20 (JC15Δ2) and depleted of AFG3L2 after inhibition of cytosolic translation with emetine. **g**, Degradation rates of DNAJC15 in (f) normalized to time point 0 h. Data are means +/− SD (n = 3). *P* values were calculated using a 2-Way ANOVA test. **h**, Schematic representation of the OMA1- mediated DNAJC15 processing leading to a degradation by the m-AAA protease.

DNAJC15 has been identified as a short-lived mitochondrial protein ^28,29^ and we have recently observed the accumulation of DNAJC15 in HeLa cells lacking the m-AAA protease subunit AFG3L2, raising the possibility that the m-AAA protease mediates proteolysis of DNAJC15 ^28,29^. Cycloheximide (CHX)-chase experiments confirmed the rapid turnover of DNAJC15, which was completely halted in cells lacking both m-AAA protease subunits, AFG3L2 and SPG7 (Fig. 1b, c; Extended Data Fig. 1b, Supplementary Table 2). We noted that DNAJC15 accumulated in a shorter form in these cells (Fig. 1b). Analysis of the neo-amino-terminal proteome of m-AAA protease-deficient cells revealed accumulation of the amino-terminal peptide 20-35 of DNAJC15 (Fig. 1a, Supplementary Table 3), suggesting that OMA1 mediates cleavage of full-length DNAJC15 (hereafter termed L-DNAJC15) after amino acid 19 into a shorter form (hereafter termed S-DNAJC15) that is degraded by the m-AAA protease. Consistently, L-DNAJC15 accumulated in *AFG3L2*^−/−^*SPG7*^−/−^ cells upon depletion of OMA1 (Fig. 1b, c).

To confirm these findings, we generated *DNAJC15^−/−^*HeLa cell lines expressing DNAJC15, S-DNAJC15 (lacking amino acids 2-19) or a DNAJC15 variant lacking amino acids 19 and 20 (hereafter referred to as DNAJC15Δ2) (Extended Data Fig. 1c). Experiments in *DNAJC15*^−/−^ cells expressing DNAJC15 variants lacking 2, 6, or 10 amino acids at the OMA1 cleavage site had shown that deletion of amino acid 19 and 20 completely abrogated OMA1 cleavage, even after membrane hyperpolarization, which is known to activate OMA1 (Extended Data Fig. 1d, e). S-DNAJC15 was targeted to mitochondria and stabilized upon depletion of the m-AAA protease in cells expressing DNAJC15 or S-DNAJC15 (Fig. 1d, e). Remarkably, impaired OMA1 cleavage led to a significant stabilization of DNAJC15Δ2, which accumulated in WT and m-AAA protease-deficient cells (Fig. 1f, g).

We conclude from these experiments that cleavage of DNAJC15 by OMA1 after amino acid 19 leads to the destabilization of DNAJC15 and rapid proteolysis by the m-AAA protease.

### Genetic interactions link DNAJC15 to mitochondrial protein import

Complementation studies in yeast have shown functional conservation of DNAJC15 with its yeast ortholog Pam18 (Tim14) ^27,30^, suggesting that proteolysis of DNAJC15 by OMA1 and the m-AAA protease regulate mitochondrial protein import. While Pam18 (Tim14) is essential for cell growth and mitochondrial protein import in yeast ^31–33^, the basal characterization of *DNAJC15^−/−^* cells did not reveal global changes in cell growth or oxygen consumption, nor did it reveal broad changes in the cellular proteome (Extended Data Fig. 2a-c). Human cells express two Pam18 paralogs, DNAJC15 and DNAJC19. Both proteins are highly homologous in the J-domain and the transmembrane region, but DNAJC15 exposes an extended amino-terminal region to the mitochondrial intermembrane space (IMS), which is cleaved by OMA1 (Extended Data Fig. 2d). We therefore hypothesized that DNAJC19 could compensate for the loss of DNAJC15. Indeed, depletion of DNAJC19 abolished cell proliferation of *DNAJC15*^−/−^ cells but did not affect the growth of WT HeLa cells (Fig. 2a). Re-expression of DNAJC15 or DNAJC15Δ2 restored cell growth (Fig. 2b-c), whereas only partial recovery was observed after expression of S-DNAJC15 (Fig. 2d-e). Thus, DNAJC15 and DNAJC19 show functional redundancy, suggesting that both play a role in mitochondrial protein import.

**Fig. 2.**
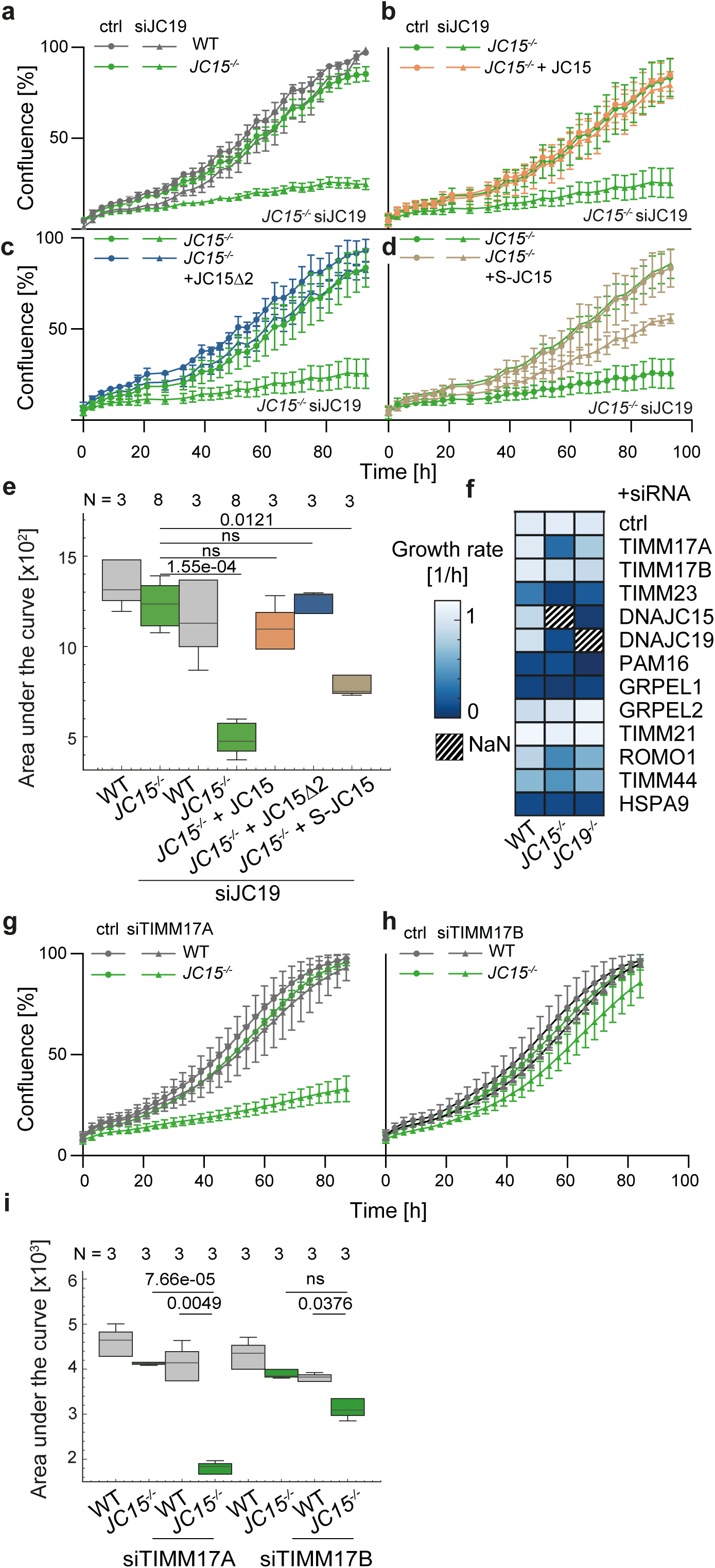
Genetic interactions of DNAJC15 with subunits of the TIM23 complex. **a**, Cell growth of wild-type (WT) and *DNAJC15^−/−^*HeLa cells after siRNA-mediated depletion of DNAJC19 (WT, n = 3; *DNAJC15*^−/−^ n = 8). Data are means +/− SD. **b**, Cell growth of WT and *DNAJC15^−/−^*HeLa cells expressing DNAJC15 (JC15) after siRNA- mediated depletion of DNAJC19 (n = 3). Data are means +/− SD. **c**, Cell growth of WT and *DNAJC15^−/−^* HeLa cells expressing non-cleavable DNAJC15 (JC15Δ2) after siRNA-mediated depletion of DNAJC19 (n = 3). Data are means +/− SD. **d**, Cell growth of WT and *DNAJC15^−/−^* HeLa cells expressing OMA1-cleaved DNAJC15 (S-JC15) after siRNA-mediated depletion of DNAJC19 (n = 3). Data are means +/− SD. **e**, Quantification of cell growth determining the area under the curve in (a-d). *P* values were calculated using a Mann-Whitney U-test. Quantile box plot show median, 25th and 75th percentiles (WT, n = 3; *DNAJC15*^−/−^, n = 8, *DNAJC15^−/−^*+ JC15, n = 3; *DNAJC15^−/−^* + JC15Δ2, n = 3; *DNAJC15^−/−^* + S-JC15, n = 3). **f**, Heatmap of growth rates of WT, *DNAJC15^−/−^* and *DNAJC19^−/−^* HeLa cells after siRNA depletion of the indicated subunits of the TIM23 complex. Data are means (n = 3). **g-i** Cell growth of WT and *DNAJC15^−/−^* HeLa cells after siRNA-mediated depletion of TIMM17A (g) or TIMM17B (h) for 72 h (n = 3). Data are means +/− SD. Quantification of cell growth monitoring the area under the curve in (g-h) are shown in (i). *P* values were calculated using a two-sided unpaired t-test. Quantile box plot show median, 25th and 75th percentiles.

We extended the analysis of the genetic interaction landscape of DNAJC15 to other components of the human TIM23 translocase ^26^. Consistent with their central role during protein translocation through TIM23 complexes, downregulation of PAM16, HSPA9, or GRPEL1 strongly inhibited cell growth in WT cells independent of the presence of DNAJC15 or DNAJC19 (Fig. 2f). Similarly, TIMM23, TIMM44, ROMO1 depletion limited the proliferation of WT cells (Fig. 2f). Cell growth was further reduced in the absence of DNAJC15, providing additional genetic support for a protein import function of DNAJC15.

Notably, depletion of TIMM17A strongly impaired the growth of *DNAJC15*^−/−^ cells, while only moderately affecting WT and *DNAJC19*^−/−^ cell growth (Fig. 2g-i). Re-expression of DNAJC15, DNAJC15Δ2 or S-DNAJC15, restored the growth of *DNAJC15*^−/−^ cells depleted of TIMM17A (Extended Data Fig. 2e-h). In contrast, DNAJC15 did not genetically interact with the TIMM17A-homologue TIMM17B (Fig. 2h, i). TIM17 proteins are an essential part of the protein translocation channel in TIM23 complexes ^34–36^. Since both human TIMM17 homologs have been found as part of distinct protein-conducting complexes ^25^, these results suggest that DNAJC15 exerts translocase-specific functions during mitochondrial protein import.

### The interactome of DNAJC15

To define the molecular environment of DNAJC15 within mitochondria, we employed immunoprecipitation coupled with chemical crosslinking to isolate untagged DNAJC15 and associated proteins. LC-MS/MS analysis revealed a significant enrichment of 185 proteins, 179 of them being mitochondrial (according to MitoCop^29^), the vast majority of which are localized to the mitochondrial matrix and IM (Fig. 3a, b, Supplementary Table 4). Consistent with previous reports ^27^, PAM16 emerged as the strongest interactor of DNAJC15 (Fig. 3c). Moreover, we identified TIMM23, TIMM44, HSPA9, TIMM50, TIMM21 and DNAJC19 as significant DNAJC15 interactors, further supporting the association of DNAJC15 with the TIM23 translocase (Fig. 3c).

**Fig. 3.**
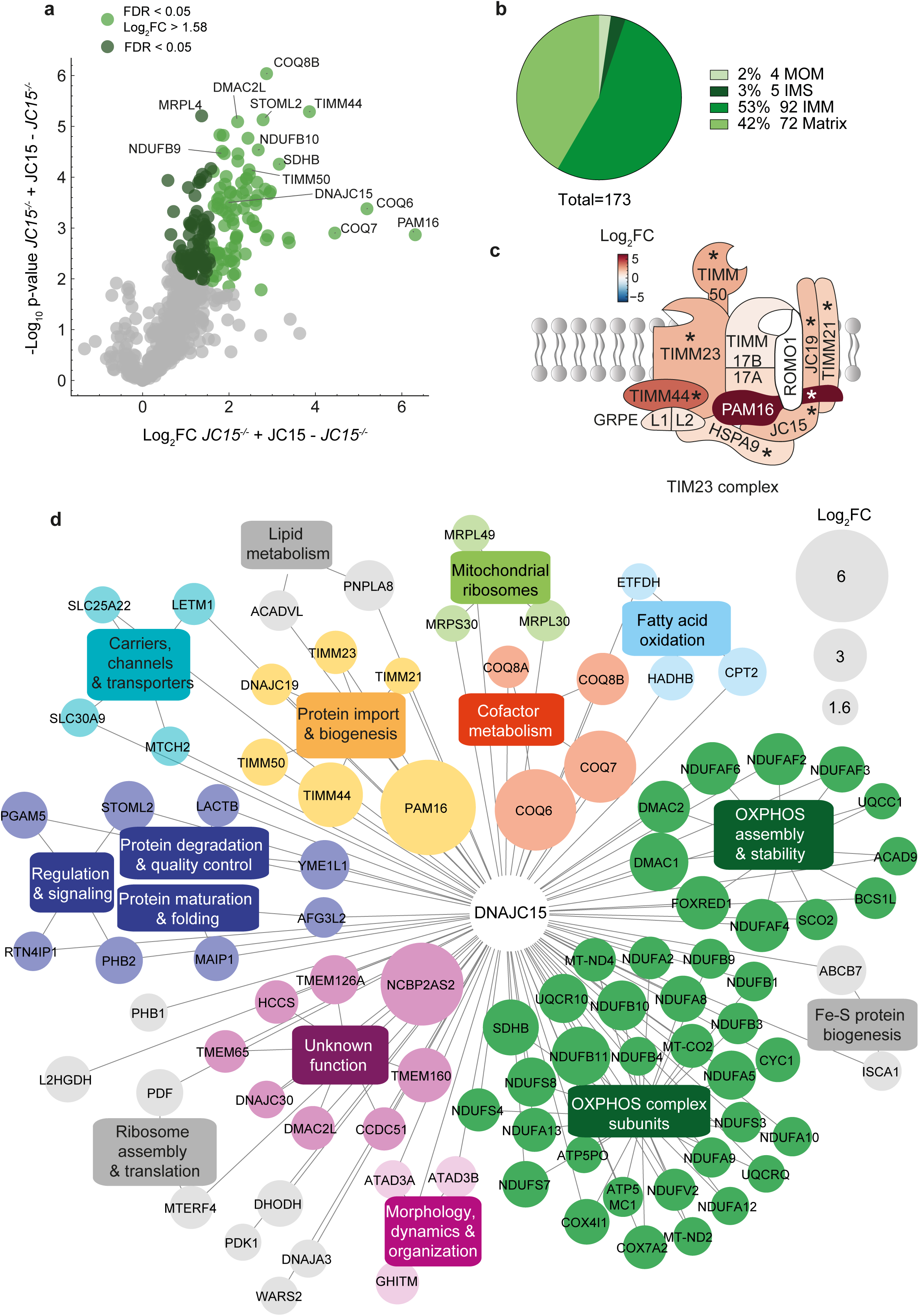
The DNAJC15 interactome. DNAJC15 immunoprecipitation in *DNAJC15^−/−^* HeLa cells expressing DNAJC15 and in *DNAJC15^−/−^* HeLa cells after chemical crosslinking (DSP, 2 mM) under non-denaturing conditions. **a**, Volcano plot showing the mitochondrial proteins interacting with DNAJC15 (according to MitoCarta 3.0). 174 significantly mitochondrial interactors are highlighted in green (FDR < 0.05, n = 4) with those that are 1.5x enriched in dark green (FDR < 0.05, log2FC < 1.58). **b**, Pie chart visualizing the known submitochondrial localization of the 173 significant mitochondrial DNAJC15 interaction partners (according to MitoCarta 3.0) (FDR < 0.05). **c**, Schematic visualization of the TIM23 complex subunits interacting with DNAJC15. The red scale indicates fold-enrichment between *DNAJC15^−/−^* cells expressing DNAJC15 and *DNAJC15^−/−^* cells (log2FC). The asterisk indicates all significantly affected proteins (FDR < 0.05, n = 4). **d**, String analysis of mitochondrial DNAJC15 interactors with a fold-enrichment higher than 1.58 (FDR < 0.05). The box size is adapted to their fold-enrichment (log2FC) in *DNAJC15^−/−^* HeLa cells expressing DNAJC15 compared to *DNAJC15^−/−^* HeLa cells. Pathways associated with only one gene name were removed.

An interaction network based on MitoCop pathways revealed close proximity of DNAJC15 to mitochondrial quality control and biogenesis factors (Fig. 3d). These include the m-AAA protease subunit AFG3L2 and its interactors MAIP1 and TMBIM5 (GHITM), the i-AAA protease YME1L as well as the CLPB chaperone and the scaffold proteins PHB1, PHB2 and STOML2 (SLP2), which are associated with AAA proteases and affect OXPHOS biogenesis (Fig. 3e) ^37–41^. Strikingly, the vast majority of proteins in the vicinity of DNAJC15 function in mitochondrial gene expression and respiratory chain biogenesis (Fig. 3d), highlighting the emerging coupling between protein translocation and OXPHOS biogenesis ^42,43^. Strikingly, the most highly enriched interactors of DNAJC15 included enzymes involved in the coenzyme Q metabolism, components involved in the respiratory complex I assembly and subunits of complex I, in particular those in the Q module and the ND4 module (Fig. 3d).

### DNAJC15 controls mitochondrial protein import specificity

Since the genetic and protein interaction landscapes linked the function of DNAJC15 to mitochondrial protein import, we directly monitored mitochondrial protein biogenesis using stable isotope labelling with amino acids in cell culture (SILAC) combined with cellular fractionation. We labelled WT HeLa cells with different stable isotopes of lysine and arginine prior to transfection with DNAJC15-specific, DNAJC19-specific or scrambled siRNA (Fig. 4a). This allowed pooling of the three samples prior to cell fractionation to reduce experimental variation. We then isolated the mitochondrial fraction by differential centrifugation and removed potential non-imported mitochondrial preproteins with exogenously added protease. The total cell fraction and mitochondrial fractions with and without protease treatment were subjected to LC-MS/MS analysis in four independent biological replicates (Fig. 4a). In total, 8342 proteins were quantified, covering 952 (83.8%) MitoCarta 3.0 annotated proteins. The SILAC ratios of individual proteins between depleted and non-depleted cells reflected changes in their steady-state levels under the different experimental conditions (Supplementary Table 5).

**Fig. 4.**
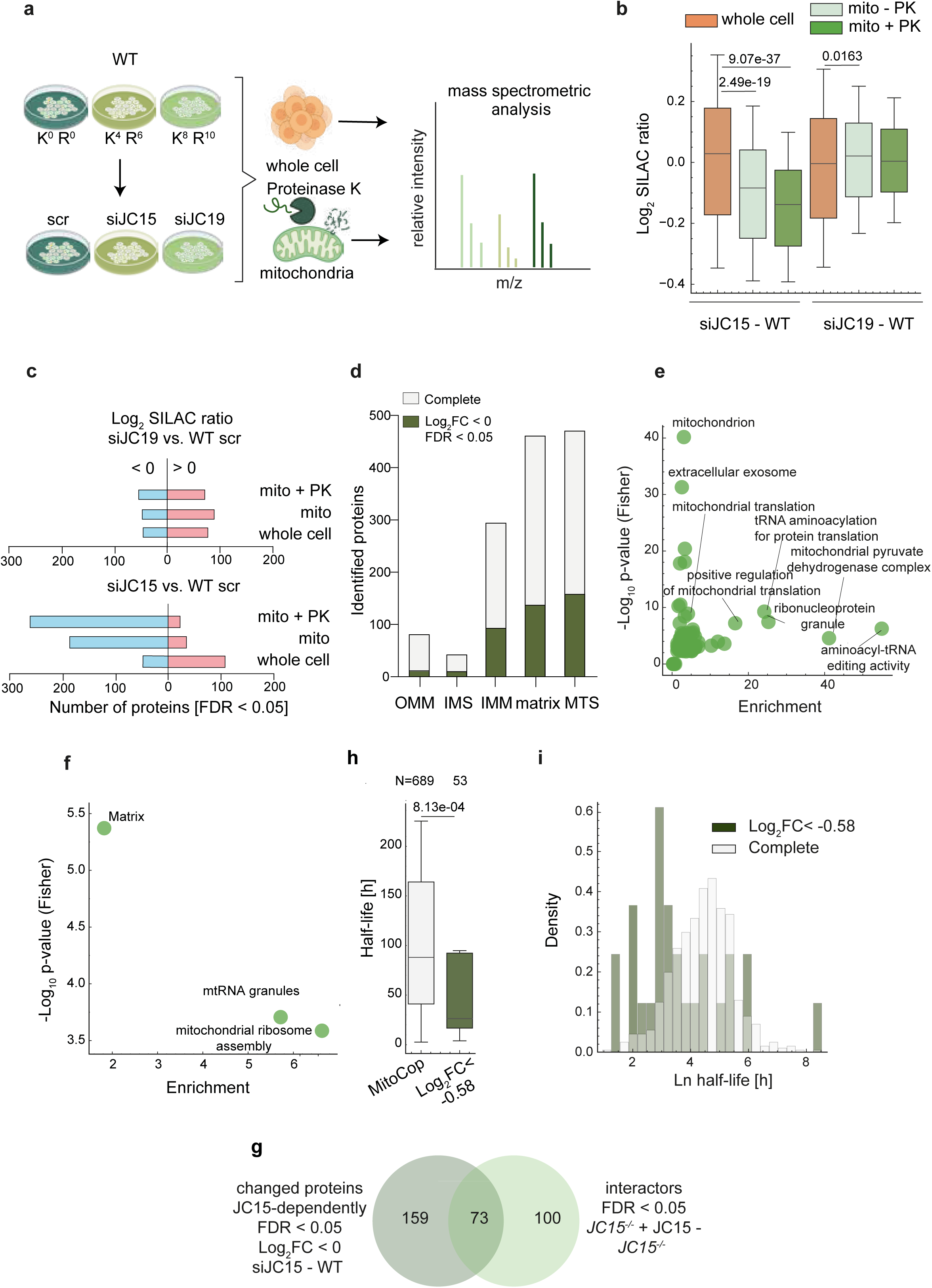
Mitochondrial import defects after depletion of DNAJC15. **a**, Wild-type HeLa cells (WT) were labelled with different radioisotopes using stable isotope labelling with amino acids in cell culture (SILAC) for a minimum of five splits, to adapt cellular pools of stable isotope labeled lysine and arginine. Differently labelled cells were depleted of DNAJC19 (JC19) and DNAJC15 (JC15) for 48 h and an equal number of cells corresponding to the three different conditions were pooled prior to subcellular fractionation, to minimize experimental variation. Samples were analyzed by LC-MS/MS (n = 5). **b**, Boxplot visualizing the distribution of mitochondrial proteins (MitoCarta 3.0) in the whole cell and in the mitochondrial fractions, which were treated with Proteinase K (PK) when indicated. Mann-Whitney U-test was performed. Quantile box plot show median, 25th and 75th percentiles were adapted to 0.5x fold (n = 5). **c**, Mitochondrial proteins (MitoCarta 3.0) significantly affected in the different fractions (FDR < 0.05) after depletion of DNAJC19 (siJC19, upper panel) and DNAJC15 (siJC15; lower panel), separated according to their SILAC ratio compared to WT (SILAC ratio > 0 for upregulation and SILAC ratio < 0 for downregulation). **d**, Fraction of significantly changed mitochondrial proteins (FDR < 0.05) within all identified proteins of various mitochondrial subcompartments and among proteins harboring a mitochondrial targeting sequence (MTS). **e**, Gene ontology enrichment analysis of all significant protein groups in the mitochondrial, Proteinase K-treated fraction after DNAJC15 depletion relative to WT (FDR < 0.02). **f**, Mitochondrial pathways enrichment analysis (MitoCarta 3.0) of significantly changed mitochondrial proteins after DNAJC15 depletion relative to WT (FDR < 0.02). **g**, Venn-diagram visualizing the overlap of significantly changed mitochondrial proteins after DNAJC15 depletion in HeLa cells (mitochondrial fraction treated with Proteinase K; Fig. 4d, n = 5) and DNAJC15 interactors (Fig. 3c) (FDR < 0.05, n = 4). **h**, Boxplot visualizing the half-life distribution of the mostly affected proteins (Log2FC < -0.58) compared to MitoCoP-annotated proteins. Mann-Whitney U-test was performed (n = 5). Half-life was extracted Morgenstern et al. ^29^. **i**, Density plot visualizing the Ln half-life distribution of the mostly affected proteins (Log2FC < -0.58) (n = 5). Half-life was extracted from Morgenstern *et al.* ^29^.

Depletion of DNAJC15 or DNAJC19 only moderately affected the distribution of mitochondrial proteins in the total cell fraction (Fig. 4b, c). We observed an only moderate increase of 110 and 77 mitochondrial proteins in cells depleted of DNAJC15 or DNAJC19, respectively (Fig. 4b, c). Thus, mitochondrial proteins are synthesized at similar rates regardless of the presence or absence of DNAJC15 or DNAJC19. Strikingly, however, 188 MitoCarta 3.0 annotated proteins were reduced in the untreated mitochondrial fraction after DNAJC15 depletion, along with a significant systematic down-shift in the distribution of mitochondrial proteins in DNAJC15-depleted cells relative to wild-type cells (Fig. 4b, c). After protease treatment of the mitochondrial fraction the steady-state level of a further 75 proteins were significantly reduced, indicating that these proteins were not imported but remained associated with the outer mitochondrial membrane (OM) in DNAJC15-depleted mitochondria (Fig. 4b, c). In contrast, the depletion of DNAJC19 did not significantly reduce stead-state levels of mitochondrial proteins (Fig. 4c).

The loss of DNAJC15 impaired mainly the import of proteins, which are localized to the matrix and IM, and of a few IMS and OM localized proteins (Fig. 4d). Many of matrix and IM proteins are targeted to mitochondria by an amino-terminal mitochondrial targeting sequence (MTS). Consistently, we observed the downregulation of MTS peptides in the mitochondrial fraction of DNAJC15-depleted cells (Fig. 4d). Gene enrichment analysis of significantly changed proteins revealed broad effects on mitochondrial proteins with diverse functions in mitochondrial gene expression, protein biogenesis and metabolism (Fig. 4e). Among the downregulated mitochondrial proteins, proteins associated with mitochondrial ribosome assembly and RNA granules were most affected as shown by a MitoPathway analysis (Fig. 4f). These results show that DNAJC15 supports the mitochondrial import of many matrix and IM proteins with OXPHOS-related functions. Consistently, 73 proteins, which were present at reduced levels after DNAJC15 depletion, are also significantly enriched in the interactome of DNAJC15 and may bind to DNAJC15 during membrane translocation into mitochondria (Fig. 4g).

The mitochondrial proteins that were significantly decreased after DNAJC15 have a decreased half-life when compared to the median stability of the mitochondrial proteome (Fig. 4h, i) ^29^, while we did not observe differences in other parameters, such as protein abundance, protein length or MTS length and score (Extended Data Fig. 3 a-d). It is therefore conceivable that, along with differences in import specificity, the high turnover of mitochondrial proteins may also contribute to the observed proteomic changes after impairment of protein import in DNAJC15-depleted mitochondria.

### DNAJC15 cooperates with TIMM17A in protein import

Previous biochemical experiments suggested the existence of distinct TIM23 complexes in the IM containing either TIMM17A or TIMM17B ^25^. We therefore depleted TIMM17A or TIMM17B from WT and *DNAJC15^−/−^*HeLa cells and analysed the mitochondrial proteome by LC-MS/MS (Fig. 5a, Supplementary Table 6). 204 and 50 mitochondrial proteins were significantly reduced after depletion of TIMM17A and TIMM17B, respectively (Fig. 5b). Notably, depletion of TIMM17A in *DNAJC15*^−/−^ cells decreased the steady-state levels of an additional 135 mitochondrial proteins, many of them linked to the assembly of OXPHOS complexes and mitochondrial ribosomes, leading to a significant shift in mitochondrial mass (Fig. 5b; Extended Data Fig. 3e-h). TIMM17B depletion in *DNAJC15*^−/−^ cells reduced the steady-state levels of further 73 proteins (Fig. 5b; Extended Data Fig. 3e-h). A two-way ANOVA of the data set identified 42 mitochondrial proteins with significantly changed steady-state levels and revealed a significant downregulation of many mitochondrial ribosome and OXPHOS-related proteins only in cells lacking both TIMM17A and DNAJC15 but not in cells lacking TIMM17B and DNAJC15 (Fig. 5c; Extended Data Fig. 3f). These experiments are consistent with the genetic interaction of DNAJC15 with TIMM17A but not TIMM17B and suggest that DNAJC15 cooperates with TIMM17A-containing complexes in the mitochondrial import of OXPHOS-related proteins into the matrix and IM.

**Fig. 5.**
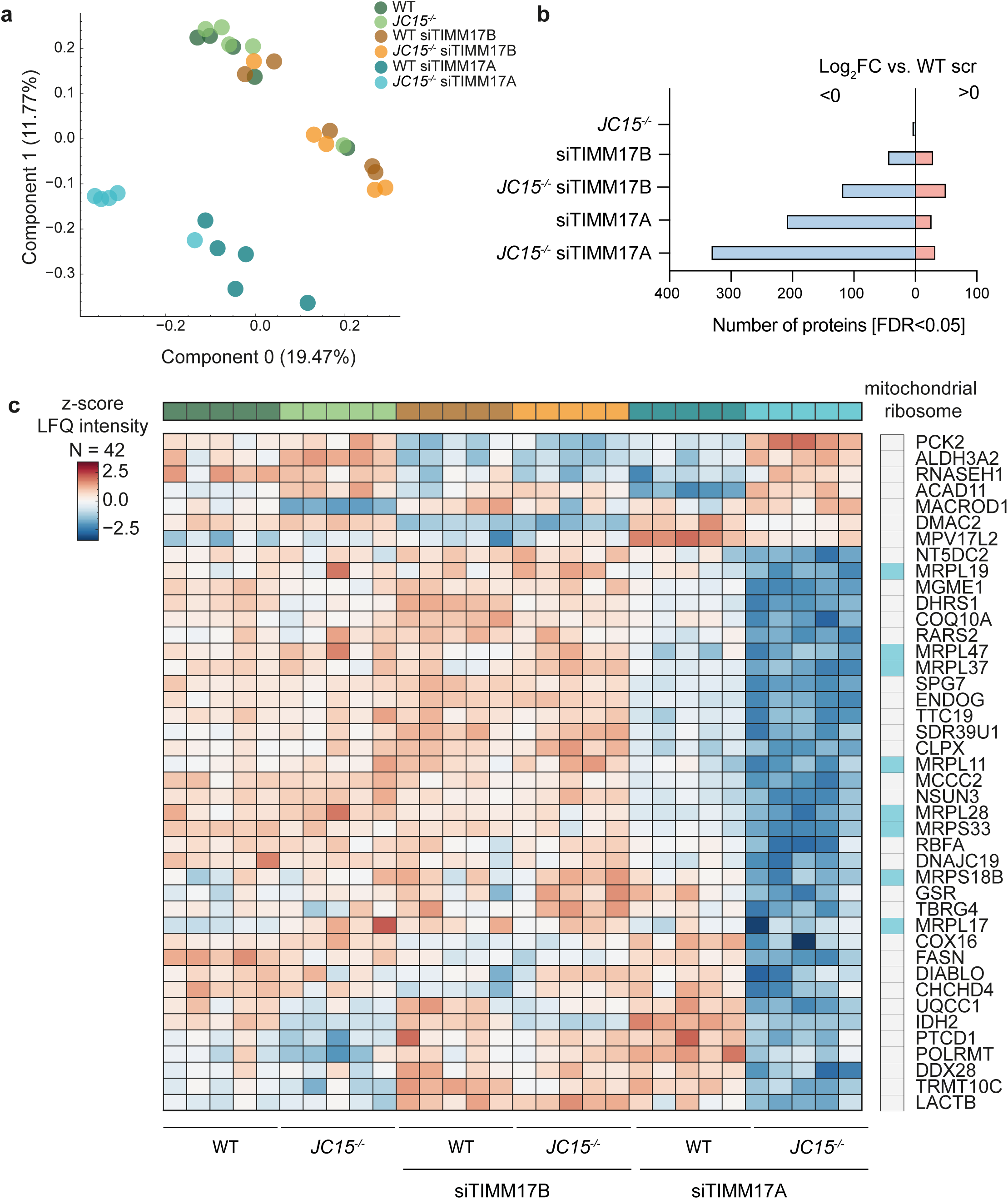
Cooperation of DNAJC15 with TIMM17A for the mitochondrial import of OXPHOS-related proteins. **a**, Principal component analysis (PCA) of cellular proteomes of wild-type (WT) and *DNAJC15^−/−^*HeLa cells after siRNA-mediated depletion of TIMM17A and TIMM17B for 72h. (n = 5). **b**, Mitochondrial proteins significantly affected by the different treatments (FDR < 0.05), separated in up-, (log2FC > 0) and downregulated (log2FC < 0) proteins. **c**, Heat-map of the two-way ANOVA significant protein groups (-log10 p-value > 2.9). Proteins corresponding to the significantly enriched gene ontology term ‘mitochondrial ribosome’ are highlighted. Dataset was filtered for minimum required values of more than 3 (n = 5).

### DNAJC15 preserves mitochondrial OXPHOS activity

Given the profound effect of DNAJC15 on the steady-state levels of numerous proteins involved in mitochondrial gene expression and OXPHOS biogenesis, we determined cellular oxygen consumption rates in isolated mitochondria depleted of DNAJC15. Consistent with previous findings ^25^, loss of DNAJC15 significantly impaired complex I activity for both phosphorylation-coupled und uncoupled states (Fig. 6a). Similarly, we observed moderately reduced complex II activity in DNAJC15-depleted mitochondria (Fig. 6b). It should be noted that the activities of respiratory complexes were not affected in the absence of DNAJC15 at the cellular level (Fig. 6c, Extended Data Fig. 2b), probably due to the moderately increased mitochondrial mass in the cells indicated by our proteomic analysis (Fig. 4c). We conclude from these experiments that DNAJC15 mediates the import of many matrix and IM proteins, whose function is linked to mitochondrial gene expression and OXPHOS biogenesis.

**Fig. 6.**
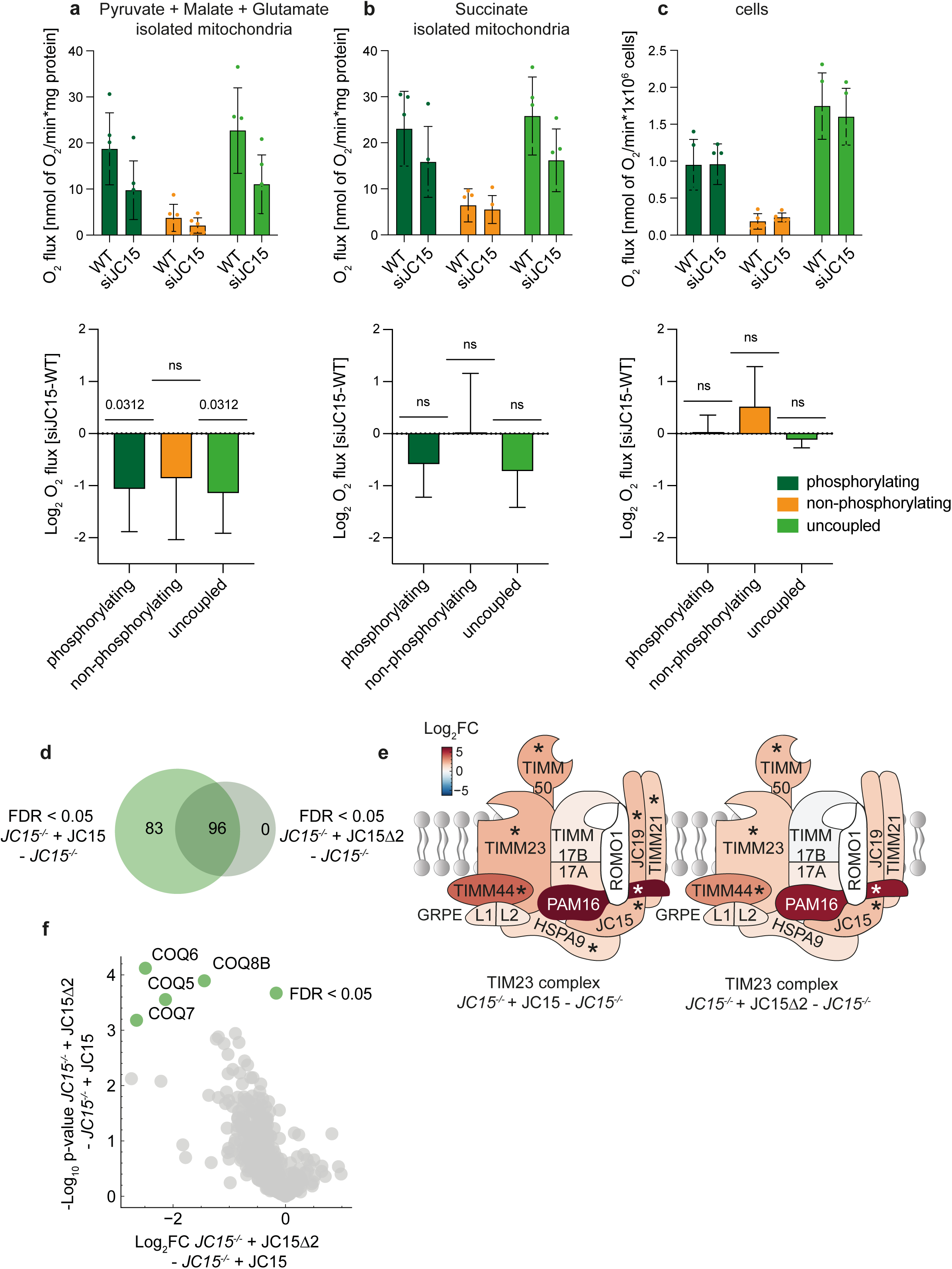
Impaired mitochondrial import after loss of DNAJC15 limits OXHOS activity. Oxygen flux was measured in intact mitochondria isolated from DNAJC15-depleted HeLa cells and wild-type (WT) cells (a, b) or in cells (c) using an Oroboros respirometer. **a**, Oxygen flux in the presence of mitochondrial complex I substrates (pyruvate 10 mM, glutamate 5 mM, malate 5 mM) under phosphorylating conditions (ADP+Pi), non-phosphorylating conditions (oligomycin, 10 µM), and after uncoupling with CCCP. Statistical significance in the log2 fold-changes in the oxygen flux between DNAJC15-depleted cells and WT cells (lower panel) was assessed using the Wilcoxon t-test. Data are means +/− SD (n = 5). **b**, Oxygen flux in the presence of mitochondrial complex II substrate (succinate 10 mM) and rotenone (140 nM) under phosphorylating and non-phosphorylating conditions and after uncoupling as in (a) (n = 4). Data are means +/− SD (n = 4). **c**, Oxygen flux in cells under phosphorylating and non-phosphorylating conditions and after uncoupling and statistical evaluation as in (a). Data are means +/− SD (n = 4). **d**, Venn diagram visualizing the overlap of the significantly identified interactors of DNAJC15 and DNAJC15Δ2 (FDR < 0.05). **e**, Schematic visualization of the TIM23 complex subunits interacting with DNAJC15. The red scale indicates fold-enrichment between *DNAJC15^−/−^* cells expressing DNAJC15 or DNAJC15Δ2 relative to *DNAJC15^−/−^*cells (log2FC). The asterisk indicates all significantly affected proteins (FDR < 0.05, n = 4). **f**, Volcano plot showing the mitochondrial proteins losing interaction with DNAJC15 after mutating of the OMA1-cleavage site (according to MitoCarta 3.0). Four significantly mitochondrial interactors are highlighted in green (FDR < 0.05; n = 4).

The binding of DNAJC15 to OXPHOS-related proteins is consistent with a role of DNAJC15 for their import and rationalizes the impaired mitochondrial respiration after depleting DNAJC15 (Fig. 3d). However, this interaction analysis does not distinguish between binding partners of DNAJC15 and an association of DNAJC15 with newly imported polypeptides during or after completion of translocation. Indeed, the yeast DNAJC15 homologue Pam18 has dual functions in protein translocation and assembly of respiratory chain complexes ^42^. As the amino-terminal region of DNAJC15 that is cleaved by OMA1 is required for the association of the chaperone with TIM23 complexes ^25^, we investigated how OMA1 cleavage affects the interactome of DNAJC15. However, the rapid turnover of S-DNAJC15 during chemical crosslinking precluded the direct identification of S-DNAJC15 interacting proteins (Extended Data Fig. 3i, j). We therefore compared the DNAJC15 interactome in *DNAJC15^−/−^* cells expressing DNAJC15 or non-cleavable DNAJC15Δ2. We identified 96 mitochondrial proteins as interacting partners of DNAJC15Δ2, all of which were also found in association with DNAJC15. Interacting proteins that did not depend on OMA1 cleavage, included the majority of TIM subunits, such as PAM16, TIMM44, TIMM23 and TIMM50, supporting an import function of L-DNAJC15 (Fig. 6e). Additional 83 proteins were detected exclusively in association with cleavable DNAJC15 (Fig. 6d, Supplementary Table 4). A direct comparison of the interactomes of DNAJC15 and DNAJC15Δ2 revealed that binding of coenzyme Q biosynthetic enzymes was most strongly affected and significantly reduced if OMA1 cleavage is impaired (Fig. 6f), suggesting that these proteins bind preferentially to S-DNAJC15.

These results show that OMA1 cleavage alters the interactome of DNAJC15. Whereas L-DNAJC15 supports protein translocation by TIM23 complexes, S-DNAJC15 appears to preferentially bind proteins involved in coenzyme Q synthesis, which is crucial for for OXPHOS biogenesis. This finding is reminiscent of the dual function of the yeast homologue Pam18 in protein translocation and assembly of respiratory chain complexes ^42^.

### Mitochondrial proteins mislocalised to the ER upon loss of DNAJC15 trigger an ATF6-related stress response

Since the loss of DNAJC15 reduced the steady-state levels of many mitochondrial proteins within the organelle but not at the cellular level, we performed further cell fractionation experiments to determine the localization of the non-imported mitochondrial proteins (Fig. 7a). After SILAC labelling and downregulation of DNAJC15 and DNAJC19, we isolated mitochondria by low speed centrifugation (8,000xg; fraction A; Fig. 4a) and further fractionated the supernatant by differential centrifugation, to enrich the endoplasmic reticulum (ER) and plasma membrane (40,000xg; fraction B) or other vesicular structures (100,000xg; fraction C). The remaining supernatant was designated as the cytosolic fraction (fraction D) (Fig. 7a). LC-MS/MS analysis of the cell fractions confirmed the enrichment of different organellar structures in the different fractions (Extended Data Fig. 4a-c) and showed an accumulation of 130 mitochondrial proteins in the fraction B containing the ER membrane only after depletion of DNAJC15 (Fig. 7b, c; Extended Data Fig. 4d, Supplementary Table 7). IM proteins and matrix proteins, which are imported along the TIM23^motor^ pathway ^44^, were enriched among proteins accumulating in fraction B (Extended Data Fig. 4e-g). 53 mitochondrial proteins that accumulated in fraction B were also significantly downregulated in fraction A. This is consistent with the observed protein import deficiency in the absence of DNAJC15. In contrast, we did not observe reduced steady-state levels of mitochondrial proteins in fraction A or accumulation of mitochondrial proteins in fraction B after depletion of DNAJC19 (Fig. 7d). Thus, these experiments suggest that non-imported mitochondrial preproteins accumulate at the ER specifically in cells lacking DNAJC15.

**Fig. 7.**
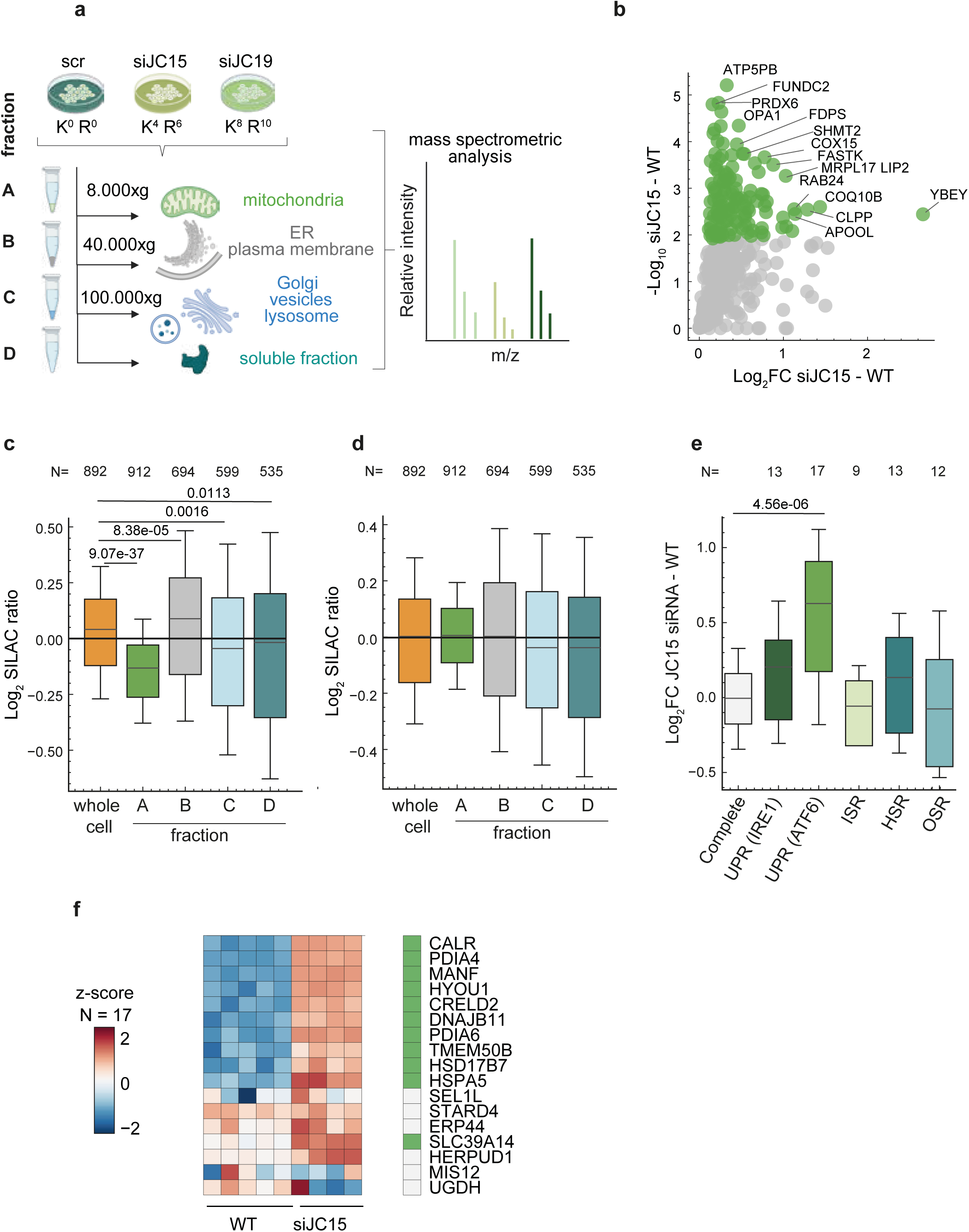
Non-imported mitochondrial proteins after DNAJC15 loss induce an unfolded protein response at the ER. **a**, Subcellular fractionation of SILAC-labelled cells depleted of DNAJC15 and DNAJC19 (see Fig. 4a) by differential centrifugation, followed by LC-MS/MS analysis. (n = 5). **b**, Volcano plot of mitochondrial proteins accumulating in the 40.000xg fraction (Significant proteins, FDR < 0.05, are labelled in green) (n = 5). The 130 most significantly proteins are labelled in green. **c**, Boxplot visualizing the distribution mitochondrial proteins (MitoCarta 3.0) in the different cellular fractions as a SILAC ratio (log2) between DNAJC15-depleted and wild-type (WT) cells (n = 5). Mann-Whitney U-test was performed. Quantile box plot show median, 25th and 75th percentiles are adapted to 0.5x fold (n = 5). N indicates the number of proteins identified in each fraction. **d**, Boxplot visualizing the distribution of mitochondrial proteins (MitoCarta 3.0) in all different fractions as a (c) after DNAJC19 depletion (n = 5). N indicates the number of proteins identified in each fraction. **e**, Gene expression analysis of wild-type (WT) HeLa cells and HeLa cells depleted of siDNAJC15 for 48 h by RNA sequencing. Box plot visualizing different transcriptionally-regulated stress responses according to gene sets regulated downstream of the following signaling pathways ^48^: the unfolded protein response (UPR) depending on IRE1 or ATF6; the integrated stress response (ISR); the heat shock response (HSR) and the oxidative stress response (OSR). Mann-Whitney U-test was performed. Quantile box plot show median, 25th and 75th percentiles (n = 5). **f**, Heat-map of ATF6-related UPR targets^48^ whose expression in DNAJC15-depleted relative to wild-type (WT) cells (log2FC) is shown. Genes whose expression was significantly changed after DNAJC15 depletion (p < 0.05) are shown in green.

The accumulation of misfolded proteins and disturbances in ER proteostasis trigger ER stress responses^45,46^. Therefore, we performed RNA sequencing under DNAJC15-depleted conditions and used a gene set profiling approach, which was developed to identify the activation of stress-responsive signaling pathways ^47,48^. These pathways included the integrated stress response (ISR), heat shock response (HSR), oxidative stress response (OSR), and the different branches of the ER unfolded protein response (UPR) mediated by IRE1 or ATF6 (Fig. 7e, Supplementary Table 8).

Loss of DNAJC15 did not induce an ISR, as indicated by the unaltered expression of the known target genes of the ISR or the ISR^mt^ (Fig. 7e; Extended Data Fig. 5a, b). OMA1, which can trigger the ISR^mt^ through DELE1 processing, was not activated in these cells nor was OPA1 processing increased (Extended Data Fig. 5c). However, DNAJC15 depleted cells exhibited the expected ISR upon oligomycin treatment, demonstrating that these cells are not refractory to ISR induction (Extended Data Fig. 5b). Similar to the ISR, the loss of DNAJC15 did not induce either the OSR or the HSF1-dependent HSR, which was observed after HSP90 inhibition and perturbation of mitochondrial proteostasis leading to UPR^mt^ ^49^. In contrast, we observed activation of the ATF6-mediated UPR, as indicated by the increased expression of 11 downstream genes (Fig. 7f), whereas the IRE1/XBP1-mediated UPR was not induced (Fig. 7e).

We conclude from these experiments that the loss of DNAJC15 impairs mitochondrial protein import and leads to the accumulation of mitochondrial preproteins at the ER, disrupting ER proteostasis and triggering an ATF6-related UPR.

## Discussion

We have identified DNAJC15 turnover as part of the OMA1-mediated mitochondrial stress response, which thus integrates changes in mitochondrial morphology and stress signalling with protein import regulation. In response to mitochondrial deficiencies, OMA1 cleavage at the amino terminus of DNAJC15 initiates its degradation by the m-AAA protease, thereby limiting the import of newly synthesized, OXPHOS-related proteins into mitochondria. We propose that this reduces OXPHOS biogenesis under stress, until mitochondrial function can be restored or damaged mitochondria are removed by mitophagy ^50^.

DNAJC15, together with its paralogue DNAJC19, supports mitochondrial protein import through TIM23 complexes as part of the import motor at the matrix side of the IM ^25^. Consistently, we identified PAM16 (MAGMAS), DNAJC19, HSPA9, TIMM23, TIMM44 and TIMM21 in the vicinity of DNAJC15. DNAJC15 binding to TIM23 complexes did not depend on OMA1 cleavage, identifying L-DNAJC15 as the import competent form. PAM16 recruits DNAJC15 (and DNAJC19) to TIM23 complexes and ensures HSPA9-dependent protein import ^25,30,51^. DNAJC15 and DNAJC19 interact genetically, suggesting at least partially redundant functions. However, in contrast to DNAJC19, loss of DNAJC15 broadly affects protein biogenesis and impairs the import of proteins with OXPHOS-related functions. Depletion of TIMM17A but not of TIMM17B aggravates this effect and further reduces the import of OXPHOS-related proteins into DNAJC15-deficient mitochondria, suggesting that DNAJC15 cooperates with TIMM17A in protein import. Indeed, previous biochemical experiments have identified different protein translocases harboring TIMM17A and TIMM17B that cooperate with DNAJC15 and DNAJC19 ^25^. Notably, in contrast to their paralogues, TIMM17A and DNAJC15 are unstable proteins ^29,52^. Our results suggest that the proteolytic removal of this protein translocase as part of the OMA1-mediated stress response allows modulation to the mitochondrial import specificity to cope with unfavourable stress conditions ^53^.

The regulation of DNAJC15 by proteolysis highlights the importance of the relative steady-state levels of components of the translocation machinery for protein import. Consistently, overexpression of PAM16, as observed in cancer, has been shown to affect the recruitment of DNAJC15 and DNAJC19 to the respective protein translocases ^30^. It is therefore conceivable that different relative expression levels of translocase subunits in different cells may explain the apparently discrepant effects of DNAJC15 depletion on OXPHOS activities in immune cells ^54–57^ (Extended Data Fig. 6).

How does DNAJC15 specifically affect the biogenesis of OXPHOS-related proteins and OXPHOS biogenesis? While TIMM17A- and TIMM17B-containing translocases may exhibit different substrate specificities, we noted overall shorter half-lives of mitochondrial proteins that accumulated in mitochondria in a DNAJC15-dependent manner. It is therefore conceivable that a generally reduced import capacity after DNAJC15 degradation and rapid proteolysis of already imported mitochondrial proteins contribute to the observed changes in the mitochondrial proteome.

Loss of DNAJC15 does not inhibit protein import into mitochondria or completely block translocation by precursor stalling, but leads to accumulation of mitochondrial preproteins at the ER. Disruption of ER proteostasis in the absence of DNAJC15 triggers an ATF6-related UPR, demonstrating coupling of mitochondrial and ER proteostasis. Similarly, perturbations in the mitochondrial lipid transport can alter the lipid composition in the ER and trigger a UPR ^58^. The accumulation of mitochondrial preproteins at the ER is reminiscent of studies in yeast suggesting that some mitochondrial preproteins encounter the ER membrane on their way to the mitochondria ^59,60^. Accordingly, the accumulation of mitochondrial preproteins at the ER upon loss of DNAJC15 during mitochondrial stress would allow a rapid recovery of mitochondrial import upon relief of the stress. While it remains to be established whether ER-associated preproteins remain competent for mitochondrial import, our findings reveal a close link in organellar proteostasis regulation and an integrated cellular stress response to mitochondrial dysfunction.

## Extended Data Figure Legends

**Extended Data Fig. 1.**
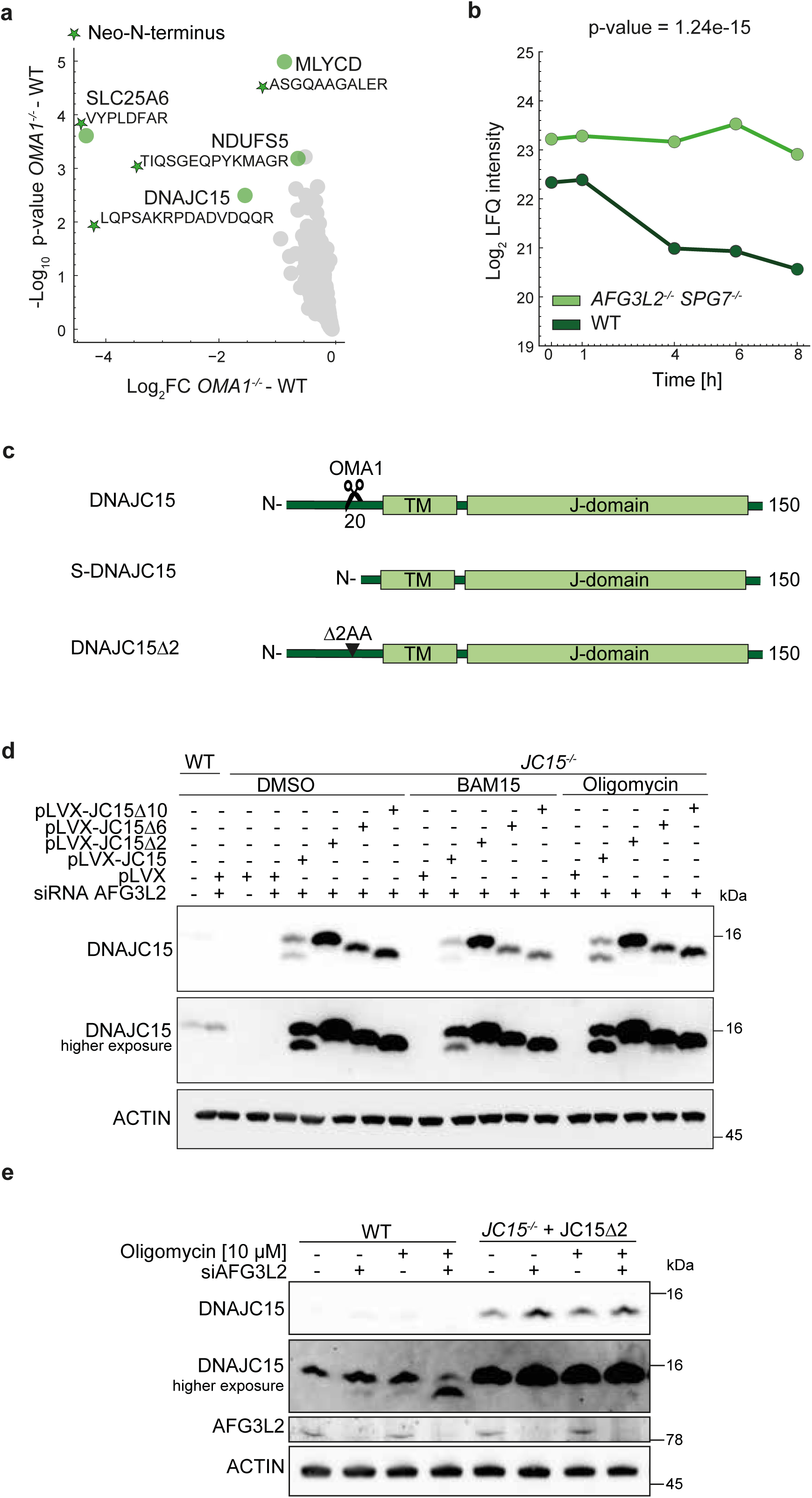
**a**, Volcano plot of the Neo-N-terminal proteome of wild-type (WT) and *OMA1^−/−^* HeLa cells, highlighting significantly affected mitochondrial proteins in green (FDR < 0.05; MitoCarta 3.0). (n = 5). **b**, Stability of DNAJC15 in WT and *AFG3L2^−/−^ SPG7^−/−^* HeLa cells (FDR < 0.05; n = 5). **c**, Schematic representation of the different DNAJC15 variants (TM = transmembrane domain). **d**, Western blot analysis of WT and *DNAJC15^−/−^* HeLa cells, which transiently express DNAJC15 or DNAJC15 variants lacking two (19/20), six (17-22) or ten (15-24) amino acids. After siRNA-mediated depletion of AFG3L2 for 72 h and treatment with oligomycin (10 µM) or Bam15 (10 µM) for 2 h, cells were lyzed and analyzed by SDS-PAGE and immunoblotting. **e**, Western blot analysis of WT and *DNAJC15^−/−^* HeLa cells stably expressing DNAJC15Δ2 following treatment with siRNA directed against AFG3L2 for 72 h and with oligomycin (10 µM) for 16 h as indicated.

**Extended Data Fig. 2.**
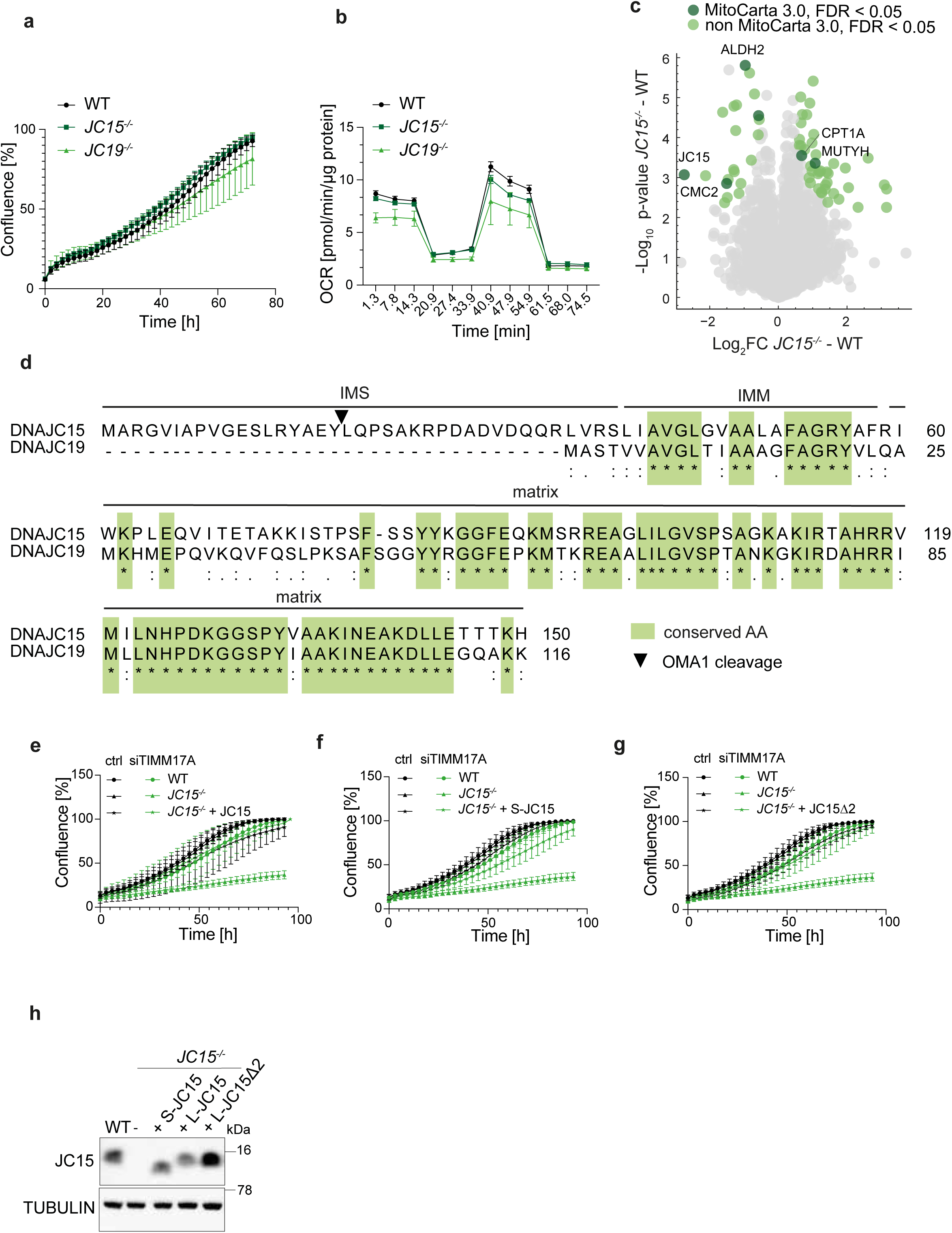
**a**, Cell growth of wild-type (WT), *DNAJC15^−/−^*, and *DNAJC19^−/−^*HeLa cells (n = 3). Data are presented as means +/− SD. **b**, Oxygen consumption rate (OCR) of WT, *DNAJC15^−/−^*, and *DNAJC19^−/−^* cells after the addition of oligomycin, FCCP, and antimycin A (n = 3). **c**, Volcano plot of cellular proteomes of wild-type and *DNAJC15^−/−^* HeLa cells. Significantly affected proteins (FDR < 0.05) are shown in green, significantly changed MitoCarta 3.0-annotated proteins in dark green (n = 5). **d**, Sequence alignment of human DNAJC15 and DNAJC19. Conserved amino acids are highlighted in green. IMS, intermembrane space; IMM, inner mitochondrial membrane. **e-g**, Growth of wild-type (WT) cells, *DNAJC15^−/−^* cells and *DNAJC15^−/−^* cells expressing (e) DNAJC15, (f) cleaved DNAJC15 (S-DNAJC15) or (g) DNAJC15Δ2, which were treated with siRNA targeting TIMM17A for 72 h (n = 3). Data are presented as means +/− SD. **h**, Expression of DNAJC15 and variants in different HeLa cell lines monitored by Western Blot.

**Extended Data Fig. 3.**
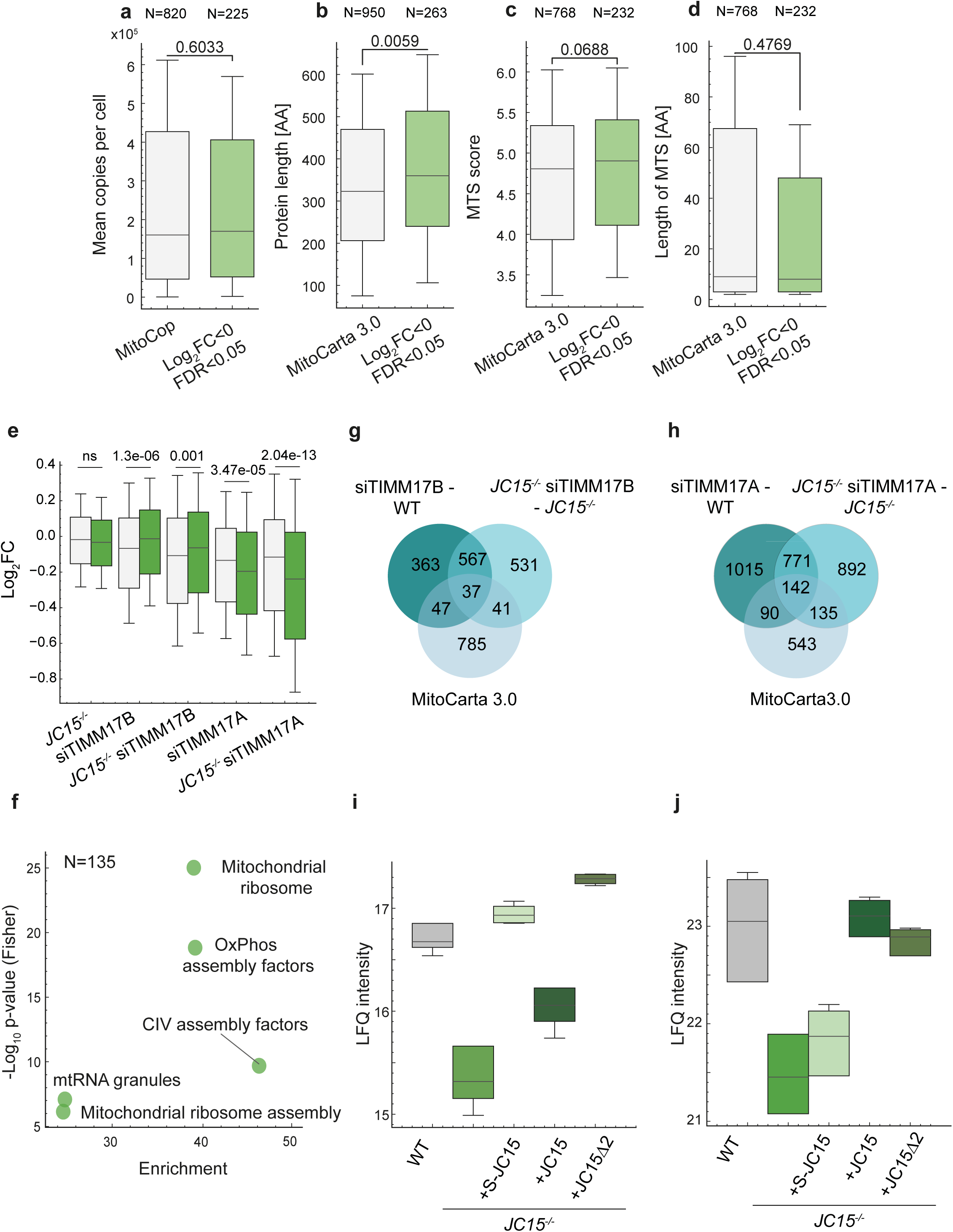
**a**, Boxplot visualizing the distribution of significantly downregulated proteins in the mitochondrial fraction (log2FC < 0, FDR < 0.05) compared to all MitoCop or MitoCarta 3.0-annotated fractions. Mann-Whitney U-test was performed. Quantile box plot show median (n = 5). The distribution indicates the mean copies per cell, extracted from Morgenstern *et al*. ^29^, in **a**, the protein length in **b**, the highest calculated MTS score in **c** and the length of the MTS with the highest MTS score in **d**, both extracted from the MTSviewer platform ^61^. **e**, Boxplot visualizing the distribution of mitochondrial proteins (MitoCarta 3.0) compared to all identified proteins in the whole cell fraction in HeLa WT and *DNAJC15^−/−^* cells, treated with siRNA targeting TIMM17A or TIMM17B. Mann-Whitney U-test was performed. Quantile box plot show median, 25th and 75th (n = 5). **f**, MitoPathways enrichment analysis of the significant mitochondrial (according to MitoCarta 3.0) proteins, that are specifically affected in *DNAJC15^−/−^* treated with siRNA targeting TIMM17A (s. Fig. 3c, FDR < 0.05). Pathways are enriched with a FDR < 0.02. **g**, Venn diagram visualizing the overlap of the significantly changed proteins of WT and *DNAJC15^−/−^*cells, treated with siRNA targeting TIMM17B, and MitoCarta 3.0-annotated protein groups (FDR < 0.05). **h**, Venn diagram visualizing the overlap of the significantly changed proteins of WT and *DNAJC15^−/−^* cells, treated with siRNA targeting TIMM17A, and MitoCarta 3.0-annotated protein groups (FDR < 0.05). **i**, Boxplot visualizing the LFQ intensity of all identified DNAJC15 peptides in the different cell lines from the input fraction after crosslinking with 2 mM DSP (n=4). **j**, Boxplot visualizing the LFQ intensity of all identified DNAJC15 peptides in the different cell lines in the elution fraction after crosslinking with 2 mM DSP (n=4).

**Extended Data Fig. 4.**
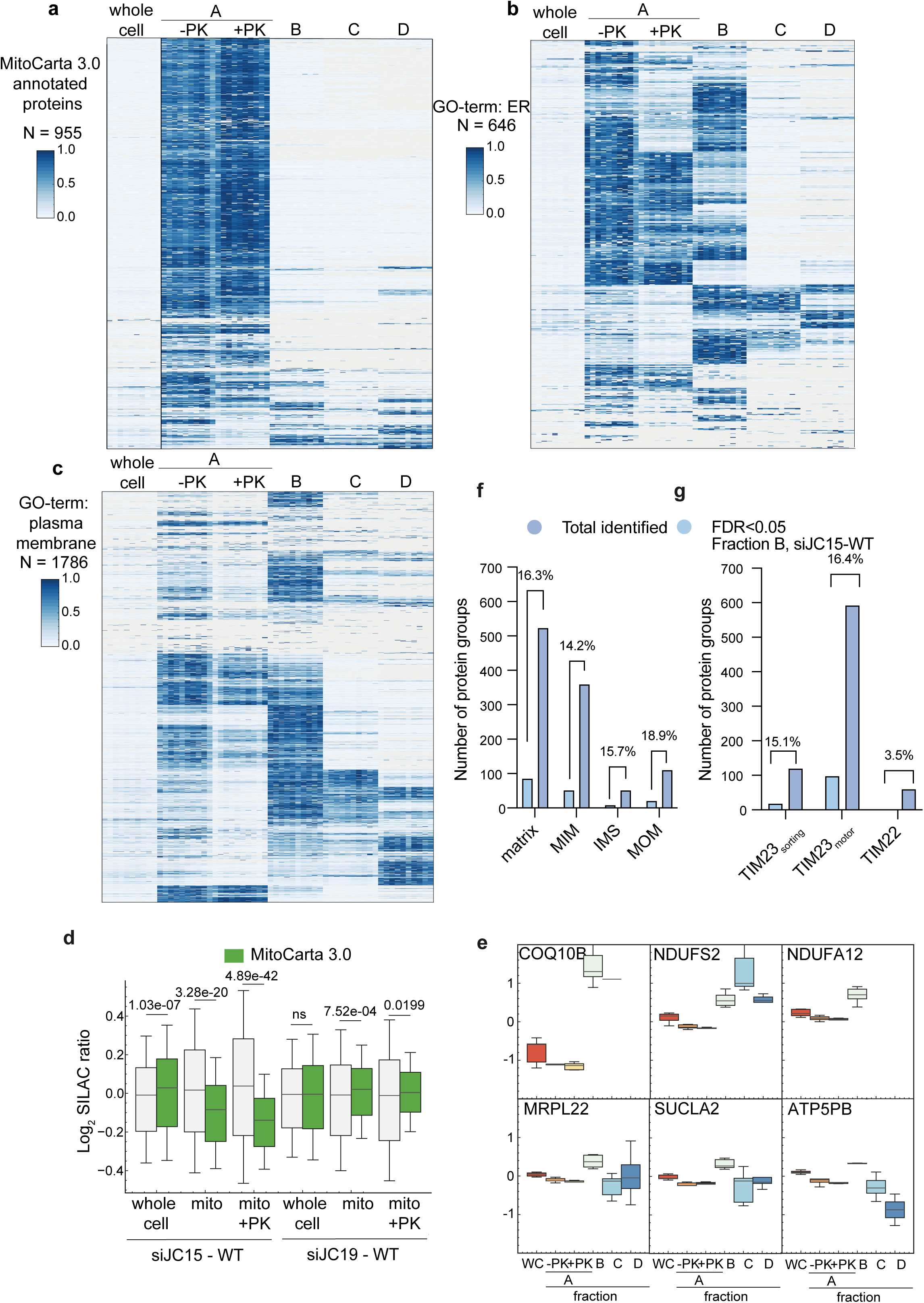
**a**, Heat map showing the intensity-based absolute quantification values of all MitoCarta 3.0 annotated proteins in all extracted fractions according to Fig. 7a (n = 5). **b**, Heat map showing the intensity-based absolute quantification values of all proteins annotated with the GO-term ‘ER’ in all extracted fractions according to Fig. 7a (n = 5). **c**, Heat map showing the intensity-based absolute quantification values of all proteins annotated with the GO-term ‘plasma membrane’ in all extracted fractions according to Fig. 7a (n = 5). **d**, Boxplot visualizing the distribution of selected mitochondrial proteins (MitoCarta 3.0) and all identified proteins in the whole cell and in the mitochondrial fractions, which were treated with Proteinase K (PK) when indicated. Mann-Whitney U-test was performed. Quantile box plot show median, 25th and 75th percentiles were adapted to 0.5x fold (n = 5). **e**, Panel of significantly downregulated proteins in the mitochondrial fraction treated with PK, that were also significantly upregulated in the 40.000xg fraction upon depletion of DNAJC15 (FDR < 0.05, n = 5). **f**, Fraction of significantly changed mitochondrial proteins (FDR < 0.05) within all identified proteins according to the submitochondrial localization of MitoCarta 3.0. **g**, Fraction of significantly changed mitochondrial proteins (FDR < 0.05) within all identified proteins according to their mitochondrial protein import in ^44^.

**Extended Data Fig. 5.**
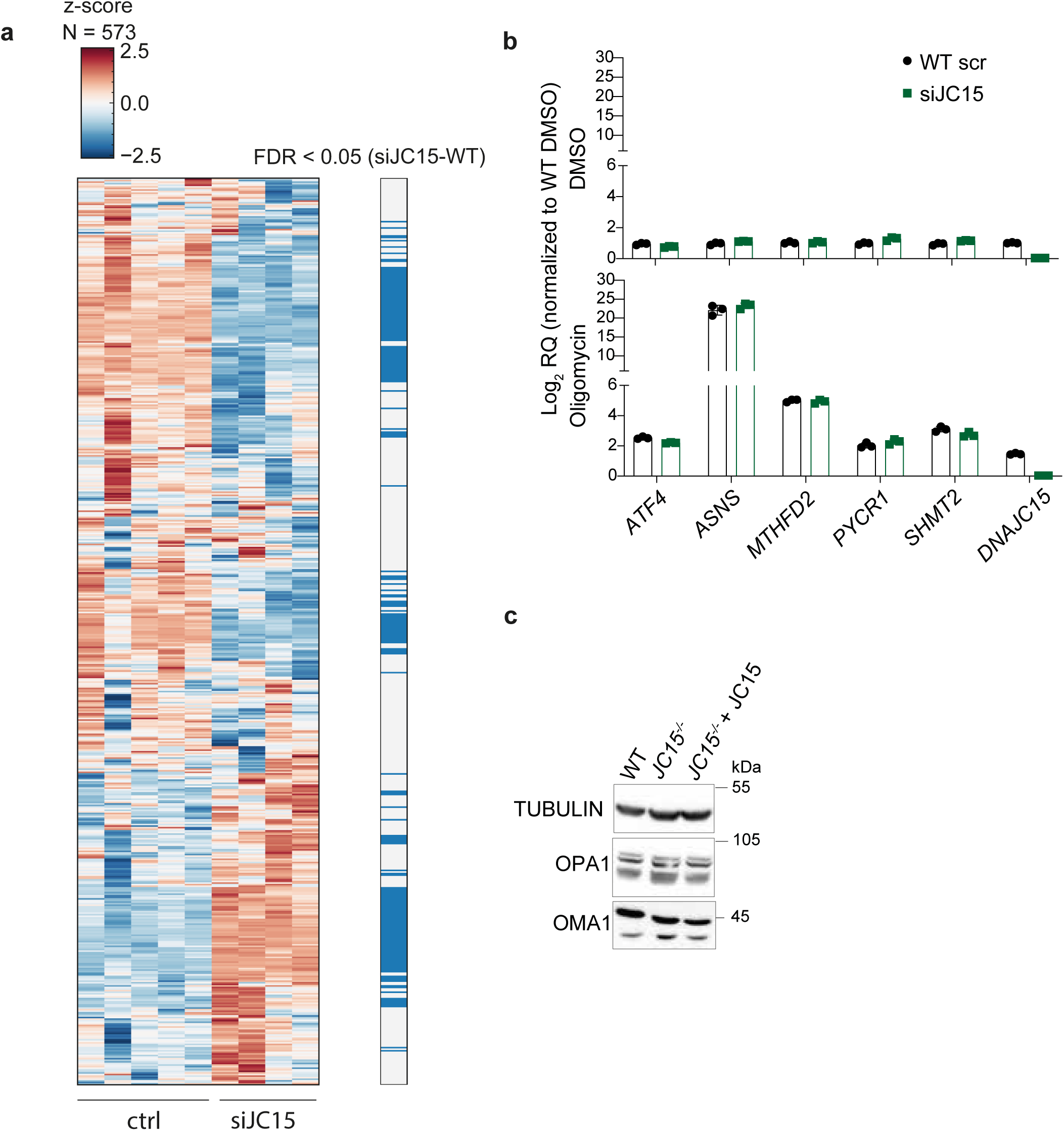
**a,** Heat map of ATF4-dependent target genes^62^ whose expression in DNAJC15-depleted relative to wild-type (WT) cells (log2FC) is shown. Genes whose expression was significantly changed after DNAJC15 depletion (p < 0.05) are shown in blue. **b**, log2 RQ values, normalized to WT cells treated with DMSO (upper panel) or oligomycin (10 µM, lower panel), for cells depleted of DNAJC15 for 48 h (n = 3). **c**, Steady-state levels of OMA1 and OPA1 in wild-type (WT), *DNAJC15^−/−^* and *DNAJC15^−/−^*cells expressing DNAJC15 analyzed by SDS-PAGE and immunoblotting.

**Extended Data Fig. 6.**
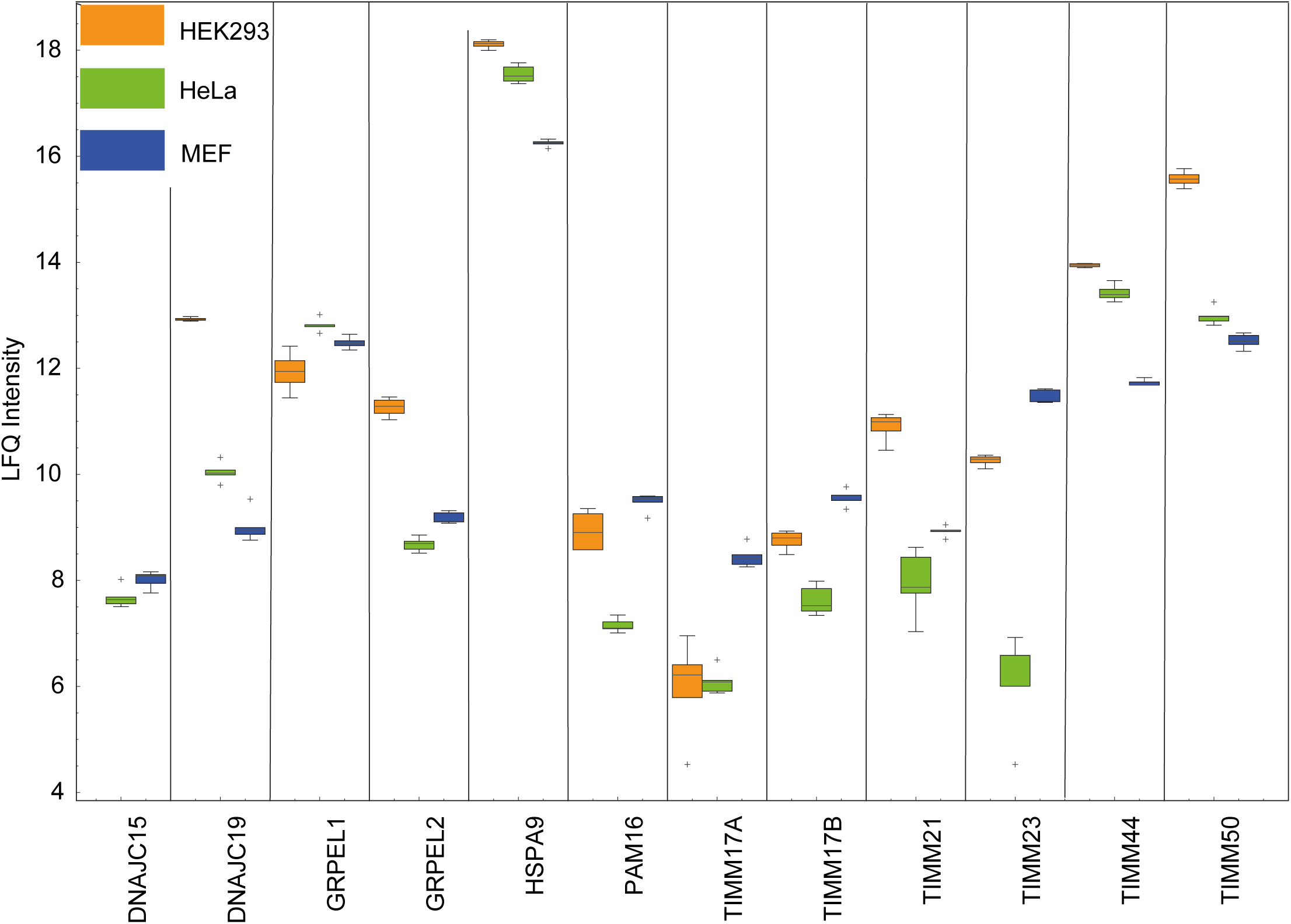
Relative expression level of the various subunits of the TIM23 complex in wild-type (WT) Human Embryonic Kidney (HEK) 293, HeLa and Mouse Embyronic Fibroblast (MEF) cell lines (n = 5).

## Supplementary Table Legends

**Supplementary Table 1.** TMT-based Neo-N terminal peptide data of WT and *OMA1^−/−^* HeLa cells.

**Supplementary Table 2.** Random mixed linear model results comparing WT and *AFG3L2^−/−^ SPG7^−/−^* HeLa cells incorporating different time points after Cycloheximide treatment. The model was calculated using the quantified peptides, utilizing genotype and time as dependent variables. The peptide variation was considered random.

**Supplementary Table 3.** TMT-based Neo N-terminal peptide detection using WT, *SPG7^−/−^*, *AFG3L2^−/−^*, and *AFG3L2^−/−^ SPG7^−/−^* knockout HeLa cells.

**Supplementary Table 4.** DNAJC15 interactome using *DNAJC15^−/−^* HeLa cells re-expressing L-JC15 (WT), s-JC15 (short) and D2-JC15 (uncleavable) DNAJC15 variants. Includes a sheet with filtered DNAJC15 interactors.

**Supplementary Table 5.** SILAC-based subcellular fractionation for the whole cell, mitochondrial and mitochondrial fraction treated with Proteinase K comparing HeLa WT treated with scrambled, DNAJC15, and DNAJC19 siRNA.

**Supplementary Table 6.** Quantified protein groups in WT and *DNAJC15^−/−^* HeLa cells using a data-independent acquisition upon knockdown of scrambled (control), TIMM17A, and TIMM17B. Contains the hierarchical clustering (Figure 5c) as a separate sheet.

**Supplementary Table 7.** SILAC-based subcellular fractionation upon siDNAJC15 and siDNAJC19 mediated knockdown for fractions B-D (40k x g, 100k x g, soluble fraction). Contains the protein group iBAQ intensities related to Extended Data Figure 4 a-c.

**Supplementary Table 8.** Transcriptomics data containing pairwise comparisons and stress gene annotations. The hierarchical clustering Figure 7f is available in a separate sheet.

## Online Methods

### Cell culture

HeLa cells were cultured in Dulbecco’s Modified Eagle’s Medium (DMEM), containing 4.5 g/L D-Glucose (Thermo Fisher, Cat# 61965-059) and 10% fetal bovine serum (Sigma, Cat# F7524). Cells were maintained at 37°C and 5% CO2 in a humidified incubator. Cell counting was performed using the viability marker trypan blue (Thermo Fisher, Cat# T10282) with the Countess automated cell counter (Thermo Fisher, Cat# AMQAF2001). Prior to each experiment, cells were seeded at equal densities and continuously monitored for Mycoplasma contamination.

### Plasmid and siRNA transfection

For transient transfection of siRNA or esiRNA, a reverse transfection was conducted with a total cell number of 4x10^5^ cells per well in a 6-well plate. The experiments were conducted 48 h or 72 h after transfection with RNAiMAXX transfection reagent (Invitrogen, Cat# 56532). The following siRNA/esiRNA targeting sequences against human proteins were utilized: siRNA AFG3L2 (SASI_HS01_0023-1520, Merck), esiRNA TIMM44 (EHU009281, Merck), esiRNA HSPA9 (EHU011841, Merck), siRNA DNAJC19 (SASI_Hs01_00055864, Sigma) siRNA DNAJC15 (SASI_Hs01_00246981, Sigma), esiRNA TIMM17B (EHU032651, Merck), esiRNA TIMM17A (EHU023551, Merck), siRNA TIMM21 (EHU031871, Merck), siRNA PAM16 (SASI_Hs01_00168312, Sigma), esiRNA TIMM23 (EHU106141, Merck), esiRNA ROMO1 (EHU224321, Merck), esiRNA TIMM50 (EHU043281, Merck), esiRNA GRPEL1 (EHU006111, Merck), esiRNA GRPEL2 (EHU065941, Merck), esiRNA OMA1 (EHU072451, Merck). For negative control, esiRNA eGFP negative control (Merck, Cat# EHUEGFP) and stealth RNAi siRNA negative control (Invitrogen, Cat# 12935300) were obtained. The efficiency of protein depletion was evaluated through immunoblotting.

### Generation of stable cell lines

Complementary cDNAs encoding human DNAJC15, S-DNAJC15 and DNAJC15Δ2 were cloned into the PiggyBac vector (PB-Cuo-MCS-IRES-GFP-EF1α-CymR-Puro) using the Q5 Site-directed mutagenesis kit (NEB, Cat# E0554S). After plasmid transfection with Genejuice Transfection Reagent (Merck, Cat# 70967) and 0.2 µg/µL of the PiggyBac Transposase Expression Vector (PB210PA-1-SBI), cells were selected with puromycin (InvivoGen, Cat# ant-pr-1) for four consecutive days after reaching full confluence. Experiments were performed with the puromycin-selected polyclonal fraction in a *DNAJC15^−/−^* HeLa cell background. Expression was induced with 8 or 15 µg/mL of 4-Isopropylbenzoic acid (Aldrich, Cat# 268402) and confirmed by immunoblotting.

### CRISPR-Cas9 gene editing

To generate *DNAJC15^−/−^* and *DNAJC19^−/−^*and *AFG3L2^−/−^ SPG7^−/−^* and *OMA1^−/−^*HeLa cells, cells were transfected with px335 plasmid (Addgene #42335), which contained guide RNAs for a nickase Cas9 deletion. The gRNAs for DNAJC15 were 5‘ACTTGCAGCCCTCGGCCAAA and 5‘GTGGTGTCATCGCTCCAGTT. The gRNAs for DNAJC19 were 5‘CGGGAAGCAGCATTAATACT and 5‘TCAGTGGTGGCTATTATAGA. The gRNAs for OMA1 were 5’CCATATAGTAAATAAGTATCAGG and 5’TGGGAGTAAATCAGTGTGACAGG. The gRNAs for AFG3L2 and SPG7 were 5’GCTGCTACCACACGCTCTTC and 5’GGTACATCAAGGCTAGCCGC, respectively. Following three consecutive transfections on subsequent days, cells were seeded as single cells in 96-well plates and examined for protein expression after a two-week growth period. Deletions were further confirmed by PCR and subsequent genomic sequencing analysis.

### IncuCyte-based cell growth assays

Cells were grown at equal densities in 96-well plates, with 5,000 cells per well. All experiments were repeated as independent biological replicates. Cell proliferation was monitored by phase microscopy imaging at 10x magnification using the IncuCyte System (Sartorius GmbH, Göttingen, Germany). To assess cell growth, the slope of log-transformed phase area was determined. The area under the curve was calculated and a U-test was performed. Significance was assessed by an unpaired two-tailed t-test.

### Protein stability assays

Cells were grown at equal densities in 6-well plates at 400,000 cells per well. Plasmid and siRNA transfection was performed for 48 h and the media was changed to complete DMEM media containing either 0.1 mg/mL Emetine (Sigma, Cat# E2375) or 100 µg/mL Cycloheximide (Sigma, Cat# C7698) for the indicated time points. Cells were collected and washed with PBS.

### Antibodies

The following antibodies were used for immunoblotting: DNAJC15 (ProteinTech, Cat# 16063-1-AP, 1:500), TUBULIN (Sigma, Cat# T6074, 1:2000), AFG3L2 (Sigma, Cat# HPA004480, 1:1000), ACTIN (Santa Cruz, Cat# SC-47778, 1:3000), TIMM17A (GeneTex, Cat# GTX108280, 1:1000), TIMM23 (BD, Cat# 611223, 1:2000), TIMM17B (ProteinTech, Cat# 11062-1-AP, 1:1000), DNAJC19 (ProteinTech, Cat# 12096-1-AP, 1:1000), SLC25A33 (OriGene, Cat# TA309042, 1:1000), TIMMDC1 (abcam, Cat# ab171978, 1:1000), SDHA (abcam, Cat# ab14715, 1:5000), OMA1 (Santa Cruz, Cat# SC-515788; 1:1,000), TOMM20 (Sigma-Aldrich, Cat# HPA011562). Corresponding species-specific HRP-coupled antibodies were used for immunoblot (Biorad; Cat# 1706515 and Cat# 1706516). The following secondary antibodies for immunochemistry were used: Anti-Mouse Alexa Fluor 488 (Thermo Fisher, Cat# A28175), Anti-Rabbit Alexa Fluor 488 (Thermo Fisher, Cat#A11008).

### SDS-PAGE and immunoblotting

Cells were harvested and washed with ice-cold PBS, before lysing with lysis buffer (50 mM (w/v) Tris-HCl pH 7.4, 1 mM (w/v) EDTA, 0.1% (w/v) SDS, 1% (w/v) Triton-X, 0.5% (w/v) sodium deoxycholate, 150 mM (w/v) NaCl) containing complete EDTA-free protease inhibitor cocktail (Roche, Cat# 64755100). Protein concentration was determined using a Bradford assay (BioRad, Cat# 5000006). Samples were boiled for 5 min at 95°C prior to SDS-PAGE. Total proteins from cells (100 µg) were separated using 14% SDS-PAGE, followed by transfer to nitrocellulose membrane and immunoblotting with the indicated antibodies. Western blot images were acquired using an Intas ChemoStar ECL Imager HR6.0 (Intas, Cat# 114801) and ChemoStar Ts software.

### Measurement of cellular respiration

Oxygen consumption rate was measured in a Seahorse Extracellular Flux Analyzer XFe96 (Agilen) according to the manufacturer’s instructions for cells grown in DMEM-GlutaMAXX containing 25 mM glucose. For each well, 3,6 x10^3^ cells were plated and incubated for 24 h at 37°C and 5% CO2 in a humidified incubator prior to measurement. Oxygen consumption rate was assessed after the addition of the corresponding OxPhos inhibitors (oligomycin – 2µM, FCCP – 0.5 µM, rotenone and antimycin A – 0.5 µM), included in the Seahorse XF Cell Mito Stress Test Kit (Agilent, Cat# 103015-100).

For respiration measurements using OROBOROS, oxygen consumption was analyzed at 37°C using isolated mitochondria (400 µg), which were suspended in 2.1 mL of respiratory buffer (MiR05 buffer: 110 mM sucrose, 0.5 mM EGTA, 20 mM Taurine, 3 mM MgCl2, 60 mM K-lactobionate, 10 mM KH2PO4, 20 mM K-HEPES; pH 7.1) or for cells (10^6^ cells) in Dulbecco’s Modified Eagle’s Medium (DMEM), containing 4.5 g/L D-glucose (Thermo Fisher, Cat# 61965-059) and 10% fetal bovine serum (Sigma, Cat# F7524). Measurements were performed with an Oxygraph-2k system (OROBOROS INSTRUMENTS, Innsbruck, Austria). Substrate-driven oxygen consumption was assessed in the presence of succinate (10 mM), pyruvate (10 mM), glutamate (5 mM), and malate (5 mM). Oxygen consumption rate was determined under three conditions: the phosphorylating state after addition of ADP (1.2 mM), the non-phosphorylating state after treatment with oligomycin (10 µM for mitochondria, 20 µM for cells), and the uncoupled state induced by the addition of carbonyl cyanide m-chlorophenylhydrazone (CCCP), which was added in incremental steps up to a concentration of 50 μM (for mitochondria) and 500 µM (for cells), to achieve maximal respiratory capacity. To ensure accurate assessment of respiratory chain-dependent oxygen consumption, antimycin A (500 nM) was added at the end of each experiment to account for residual oxygen consumption.

### Quantitative PCR

Cells were harvested and washed with ice-cold PBS prior to cell lysis using the lysis buffer provided in the RNA isolation kit (Macherey-Nagel, Cat# 740955.250). The experiment was performed according to the manufacturer’s instructions. RNA isolation was quantified using a NanoDrop spectrophotometer (Thermo Fisher, Cat# ND-ONE-W), and complementary DNA (cDNA) was synthesized using the GoScript Reverse Transcriptase kit (Promega, Cat# A2791), according to the manufacturer’s protocol, with a total RNA concentration of 50 µg/mL. Quantitative PCR was performed using SYBR Green PCR mix (Applied Biosystems, Cat# 4367659), containing 0.5 µM of each primer and 20 ng of cDNA for 40 cycles at 95°C for denaturation and 60°C for the elongation step. Hypoxanthine-guanine phosphoribosyltransferase (HPRT) was used as a housekeeping gene to normalize expression levels, which were determined using the ΔΔCt method. The sequences of the primers are provided in the material table.

### Immunoflourescence of fixed cells

HeLa cells were fixed with 4% paraformaldehyde for 10 minutes. After fixation, cells were washed with PBS for 10 and cell permeabilization was achieved by incubating cells with 0.1% Triton X-100 in PBS for 5 minutes, followed by PBS washes for each 10 minutes. Nonspecific binding sites were blocked by incubating the cells with 2% bovine serum albumin (BSA) in PBS for 30 minutes at room temperature. A mixture of primary antibodies in 2% BSA in PBS was applied and incubated overnight at 4°C. After incubation, cells were washed thoroughly with PBS for 10 minutes, repeating the wash step three times. The secondary antibody solution was prepared in 2% BSA in PBS, using antibodies conjugated to fluorophores with emission maxima of either 488 nm or 569 nm. Secondary antibodies were added and incubated in the dark for 45 minutes. After incubation, cells were washed three times with PBS for 10 minutes to remove excess secondary antibodies. For mounting, a drop of Prolong Gold Mounting Media was added to a clean glass slide. The mounted samples were allowed to dry for 1 hour at room temperature and stored at 4°C for at least 2 days.

### Neo-N-terminal proteomics

Cells were lysed and proteins were reduced (10 mM TCEP) and alkylated (20 mM CAA) in the dark for 45 min at 45°C. TMT labels (TMT10plex Label Reagent Set plus TMT11-131C, #A37725, or TMT6plex Label Reagent Set) were equilibrated to RT and solubilized in LC-MS grade acetonitrile according to manufacturer’s protocol. For 30 µg protein input, 20 µL of TMT labels were added and incubated for 1 h at 25°C on a thermomixer at 300 rpm. The reaction was stopped adding 100 mM Tris-HCL buffer to a final concentration of 20 mM. Samples were pooled to a total of 100 µg and subjected to SP3-based protein digestion. Peptides were incubated with activated NHS-magnetic beads (Pierce™ NHS-Activated Magnetic Beads, # 88826) according to the manufactor’s instructions to capture free amine groups of internal peptides (peptide N-term). The supernatant was removed, acidified and desalted using SDB-RP stage tips and resuspended in 10 mM ammonium hydroxide and 5% acetonitrile. Peptides were then separated via offline high-pH peptide fractionation. The instrumentation consisted out of a ZirconiumTM Ultra HPLC and a PAL RTC autosampler system using the binary buffer system. A) 10 mM ammonium hydroxide and B) 80% acetonitrile and 10 mM ammonium hydroxide. Peptides were separated according to their hydrophobicity using an in-house packed column (length = 40 cm, inner diameter = 175 µm, 2.7-µm beads, PoroShell, Agilent Technologies) column. The instruments communicated and were controlled using the software Chronos (Axel Semrau GmbH). The total gradient length was 40 min and in total 12 (SPG7, AFG3L2, SPG7 AFG3L2 double knockout and WT) or 36 (Oma1 knockout cell N-Termiome) fractions were collected (1/30 s) and subsequently concentrated using a SpeedVac to complete dryness.

### DSP interaction analysis

Cells were seeded at a density 15x10^6^ cells in a 15 cm cell culture dish. DNAJC15 expression was induced with 15 µg/mL of 4-isopropylbenzoic acid (Aldrich, Cat# 268402) and confirmed using immunoblotting. After 48 h of incubation in a humidified incubator, cells were washed twice with PBS. PBS containing 2 mM DSP (Thermo Fisher, Cat# 22585) was then added and the cells were incubated for 30 min at RT to allow cross-linking. For quenching, a 100 mM Tris pH 7.4 solution was added and the cells were incubated for a further 15 min at RT. Cells were then harvested, and lysed (20 mM HEPES-NaOH, 150 mM NaCl, 2 g/g digitonin (Merck, Cat# 300410)). Protein concentration was determined using Bradford reagent (BioRad, Cat# 5000006). For the subsequent immunoprecipitation, 1.4 mg of total protein was used as the input, with DNAJC15 antibody (Proteintech, Cat# 16063-1-AP) pre-bound to Protein G magnetic beads (Thermo Fisher, Cat# 10003D). The Protein G and antibody mixture was washed, 2x for 10 minutes with 0.1% Sodium Deoxycholate and 3x washed with PBS including a 10 minutes incubation step with PBS. After adding the lysate, the mixture was incubated for 120 minutes. After several washes with washing buffer (10 mM HEPES-NaOH, pH 7.5, 150 mM NaCl, 0.1% Triton-X-100), proteins were eluted by incubating the beads in Laemmli buffer (50 mM Tris-HCL, pH 6.8, 2% SDS, 10% glycerol, 0.01% bromophenol blue, 60 mM DTT) at 70°C for 10 min at 300 rpm.

### RNA sequencing data

For eukaryotic mRNA sequencing, samples were processed by Novogene. Sequencing was performed on the Illumina NovaSeq X Plus platform (PE150) using a paired-end 150 bp (PE150) read strategy. Messenger RNA was purified from total RNA using poly-T oligo-attached magnetic beads. After fragmentation, the first strand cDNA was synthesized using random hexamer primers followed by the second strand cDNA synthesis. The library was ready after end repair, A-tailing, adapter ligation, size selection, amplification, and purification. The library was checked with Qubit and real-time PCR for quantification and bioanalyzer for size distribution detection. Quantified libraries will be pooled and sequenced on Illumina platforms, according to effective library concentration and data amount.

### Mitoproteomics

HeLa cells were cultured in DMEM medium without arginine, lysine and glutamine (Silantes, Cat# 282006500), supplemented with Glutamine and 10% dialyzed FBS. Cells were adapted for seven doublings to heavy isotope media ^13^C6^15^N4 arginine (Silantes, Cat#201604102) and ^13^C6^15^N2 lysine (Silantes, Cat# 211604102), to middle isotope media ^13^C6^15^N arginine (Silantes, Cat# 201204102) and ^12^C4^15^N lysine (Silantes, Cat# 211104113), to light isotope media ^12^C^15^N arginine (Silantes, Cat# 201004102) and ^12^C^15^N lysine (Silantes, Cat# 211004102). The same medium was used for the corresponding siRNA transfection. After 48 h of incubation, cells were trypsinized, and cell counts were performed using the viability marker trypan blue (Thermo Fisher, Cat# T10282) with the Countess automated cell counter (Thermo Fisher, Cat# AMQAF2001). Equal numbers of cells were pooled, and subcellular fractionation was performed.

### Subcellular fractionation

Cell pellets were resuspended in homogenization buffer (20 mM HEPES-KOH pH 7.4, 220 mM Mannitol, 70 mM Sucrose, 1mM EGTA pH 7.4) containing complete EDTA-free protease inhibitor cocktail (Roche, Cat# 64755100). The suspension was kept on ice for a 15 min. Cell disruption was performed by applying 20 strokes in a glass mortar and PTFE pestle at 1,000 rpm using a rotating homogenizer (Schuett Biotec, Cat# 3201011). Two consecutive centrifugation steps at 600xg for 5 min were conducted to remove nuclei. Mitochondrial fractions were pelleted at 8.000xg for 10 min at 4°C. Where indicated in the figure legend, mitochondria were treated with 20 µg/mL Proteinase K (Southern Cross Science, Cat# 0219350491) for 20 min at 4°C, followed by reaction termination with 2 mM PMSF (Roche, Cat# 10837091001). For further subcellular fractionation, the supernatant of the mitochondrial-containing fraction was centrifuged at 40,000xg for 1 h, to yield the heavy microsome fraction. Additional centrifugation of the remaining supernatant at 100,000xg for 1 h allowed isolation of the light microsomal fraction. The supernatant, containing the soluble fraction was precipitated with acetone.

### Acetone precipitation

Four times the volume of ice-cold (-20°C) acetone was added to the supernatant and incubated overnight for 16 h. The samples were then centrifuged at 20,000xg for 10 min at 4°C. The pellet was washed twice with 400 µL of 80% ice-cold acetone. The pellet was dried under the fume hood for 5 min and then resuspended in 100 µL 4% SDS in 100 mM HEPES-KOH pH=8.5.

### SP3 Digestion Protocol

For total proteome analysis, 60 µL of 4% SDS in 100 mM HEPES-NaOH (pH = 8.5) was pre-warmed to 70°C and added to the cell pellet for a further 10 min incubation at 70°C on a thermomixer (shaking: 550 rpm). Protein concentration was determined using the 660 nm Protein Assay (Thermo Fisher, Cat# 22660). 20 µg of protein was subjected to tryptic digestion. For immunoprecipitations, the LDS buffer eluate was directly used. Proteins were reduced (10 mM TCEP) and alkylated (20 mM CAA) for 45 min at 45°C in the dark. Samples were subjected to an SP3-based digestion^63^. Washed SP3 beads (SP3 beads (Sera-Mag(TM) Magnetic Carboxylate Måodified Particles (Hydrophobic, GE44152105050250), Sera-Mag(TM) Magnetic Carboxylate Modified Particles (Hydrophilic, GE24152105050250) from Sigma Aldrich) were mixed equally, and 3 µL of bead slurry was added to each sample. Acetonitrile was added to a final concentration of 50% and washed twice using 70% ethanol (V = 200 µL) on a custom-made magnet. After a further acetonitrile wash (V = 200µL), 5 µL of digestion solution (10 mM HEPES-NaOH pH = 8.5 containing 0.5 µg Trypsin (Sigma, # T6567-1mg) and 0.5 µg LysC (Wako, # 129-02541) was added to each sample and incubated at 37°C overnight. Peptides were desalted on a magnet using 2 x 200 µL of acetonitrile. Peptides were eluted in 10 µL 5% DMSO in LC-MS water (Sigma Aldrich, #900682) in an ultrasonic bath for 10 min and subjected to StageTip desalting using the SDB-RPS material (Affinisep, AttractSPE®Disks SDB-RP, #SPE-Disks-Bio-DVB-47.20) ^64^. Formic acid and acetonitrile were added to a final concentration of 2.5% and 2%, respectively. Samples were stored at -20°C prior to LC-MS/MS analysis.

### Liquid Chromatography and Mass Spectrometry for Neo N-terminal proteome

LC-MS/MS instrumentation consisted out of an Easy-LC 1200 (Thermo Fisher Scientific) coupled via a nano-electrospray ionization source to an QExactive HF-x mass spectrometer (Thermo Fisher Scientific). For peptide separation an in-house packed column (inner diameter: 75 μm, length: 20 cm) was used. A binary buffer system (A: 0.1 % formic acid and B: 0.1 % formic acid in 80% acetonitrile) was applied as follows: Linear increase of buffer B from 4% to 28% within 33 min, followed by a linear increase to 55% within 5 min. The buffer B content was further ramped to 95 % within 2 min. 95 % buffer B was kept for further 3 min to wash the column. Prior each sample, the column was washed using 6 μL buffer A and the sample was loaded using 7 μL buffer A. The mass spectrometer operated in a data-dependent mode and acquired MS1 spectra at a resolution of 60000 (at 200 m/z) using a maximum injection time of 20 ms and an AGC target of 3e6. The scan range was defined from 350-1650 m/z and data type was set to profile. MS2 spectra were acquire in a Top15 mode at a 45000 resolution (at 200 m/z) using an isolation window of 0.8 m/z and a normalized collision energy of 32. The first mass was set to 110 m/z. Dynamic exclusion was enable and set to 20 s.

### Liquid Chromatography and Mass Spectrometry for data independent acquisition (whole proteome, immunoprecipitation)

The LC-MS/MS instrumentation consisted of an Easy-LC 1200 (Thermo Fisher Scientific) coupled via a nano-electrospray ionization source to an Exploris 480 mass spectrometer (Thermo Fisher Scientific, Bremen, Germany). An Aurora Frontier column (60 cm length, 1.7 µm particle diameter, 75 µm inner diameter, Ionopticks). A gradient based buffer system (A: 0.1% formic acid and B: 0.1% formic acid in 80% acetonitrile) based gradient was used at a flow rate of 185 nL/min as follows: a linear increase of buffer B from 4% to 28% within 100 min, followed by a linear increase to 40% within 10 min. The buffer B content was further increased to 50% within 4 min and then to 65% within 3 min. 95% buffer B was maintained for a further 3 min to wash the column. The RF lens amplitude was set to 45%, the capillary temperature to 275°C and the polarity to positive. MS1 profile spectra were acquired at a resolution of 30,000 (at 200 m/z) over a mass range of 450-850 m/z and an AGC target of 1 × 10^6^.

For MS/MS independent spectra acquisition, 34 equally spaced windows were acquired with an isolation m/z range of 7 Th, and the isolation windows overlapped by 1 Th. The first fixed mass was 200 m/z. The isolation center range covered a mass range of 500–740 m/z. Fragmentation spectra were acquired with a resolution of 30,000 at 200 m/z using a maximum injection time setting of ‘auto’ and stepped normalized collision energies (NCE) of 24, 28, and 30. The default charge state was set to 3 and the AGC target was 3e6 (900% - Exploris 480). MS2 spectra were acquired in centroid mode. FAIMS was activated with an inner electrode temperature of 100°C and an outer electrode temperature of 90°C. The compensation voltage was set at -45 V.

### Data Analysis RNA sequencing

rRNA transcripts were removed from the annotation file by depleting all lines with ‘rrna’ tag on it. cDNA index was build using kallisto (kallisto/0.46.1) and RSeQC/4.0.0 used to identify mapping strand: A strand was identified by having more than 60% of reads mapped to it. Cases with less than 60% of reads in each strand are defined as unstranded. After normalization of read counts by making use of the standard median-ratio for estimation of size factors, pair-wise differential gene expression was performed using DESeq2/1.24.0. After removal of genes with less then 10 overall reads log2 fold changes were shrank using approximate posterior estimation for GLM coefficients.

### Neo N terminal proteome

Raw files were analyzed using MaxQuant (1.6.7 and 1.6.12) ^65^ and the implemented Andromeda search engine. TMT 10-plex (WT and OMA1 knockout) or TMT-6plex (SPG7, L2, SPG7 AFG3L2 knockout) was set as a quantification setting. ArgC with semi-specificity (free N-terminus) was used. MS2 spectra were correlated against the Uniprot human reference proteome (UP000005640, number sequences: 21K, downloaded: 2022.08). Match-between runs algorithm was enabled. To identify novel Neo-N termini, the peptides.txt file of the MaxQuant output folder was utilized and TMT intensities between conditions were compared using a two-sided t-test. Raw files were analyzed using MaxQuant (1.6.4) ^65^ and the implemented Andromeda search engine. TMT-10plex was set as a quantification setting. ArgC with semi-specificity (free N-terminus) was used. MS2 spectra were correlated against the Uniprot human reference proteome. Match-between runs algorithm was enabled. To identify novel Neo-N termini, the peptides.txt file of the MaxQuant output folder was utilized and TMT intensities between conditions were compared using a two-sided t-test in the Perseus software.

### Whole proteome and crosslinking interaction studies

The Spectronaut (18.7.240325.55695) directDIA+ (Deep) analysis tool was used to correlate the acquired MS2 spectra in data-independent mode with the Uniprot reference human proteome (UP000005640, number sequences: 21K, downloaded: 2022-08). The MaxLFQ algorithm was used. The precursor, peptide and protein q-value cutoff were 0.01 in the Pulsar search. A total number of two missed cleavages of 2 were allowed, and the minimal peptide length was seven amino acids. Acetyl (Protein N-term), and Oxidation (M) were defined as variable modifications, and Carbamidomethyl at cysteines was set as a fixed modification. The mean precursor and mean peptide quantities were used for agglomeration to protein quantities. The log2 protein group LFQ intensities were used for a pairwise comparison using a two-tailed unpaired t-test followed by a permutation-based FDR calculation (FDR < 0.05 is considered significantly different, s0 = 0.1, number of permutations = 500). The experiment was performed with 5 (whole proteome) and 4 (interaction studies) biologically independent replicates.

### SILAC-based subcellular fractionation

The Spectronaut (18.7.240325.55695) directDIA tool was used to analyze the acquired raw files. channel 2 (Arg6, Lys4) and channel 3 (Arg10, Lys8) were defined. The option ‘exclude interference’ was enabled to exclude fragments with the same mass of different SILAC channels (e.g. co isolated b-ions). Otherwise, the default settings were utilized. The SILAC protein group ratio was then calculated by aggregating the precursor data to the mean of the log2 SILAC ratio at the elution group level and further calculating the median of the peptides log2 SILAC ratio to obtain the protein group H/L, H/M and M/L ratios. The ratio distributions were then shifted to a median of 0. Significantly different protein groups were identified using a one-sample t-test on the log2 protein group SILAC ratio. The p-values were adjusted using the Benjamini-Hochberg correction (adj. p-value < 0.05) in the Instant Clue software.

### Quantification and statistical analysis

Densitometry data were generated by Fiji for Western blot quantification. Representative images from at least three independent experiments are shown. Graphs were generated using Prism (GraphPad). Volcano plots and heat maps were generated using InstantClue (Nolte, 2018 #468). Error bars represent the standard deviation of the mean. Statistical significance of the data was assessed by using the Student’s t-test or the Whitney-Mann U-test, comparing control and test conditions, as described in the figure legend. Data was visualized in Instant Clue ^66^.

## Notes

### Competing Interest Statement

The authors have declared no competing interest.

